# Developing a Multimodal miR-15a Mimic to Overcome PARP Inhibitor Resistance in Epithelial Ovarian Cancer

**DOI:** 10.64898/2026.04.20.719456

**Authors:** Amartya Pal, Anushka Ojha, Hersh Bendale, Lei Chen, Iwao Ojima, Jingfang Ju

## Abstract

Epithelial ovarian cancer (EOC) is characterized by high relapse rates and the development of drug resistance, driven by adaptive DNA repair and survival pathways. Here, we develop a multimodal, chemically engineered miRNA therapeutic, MTX-5-FU-Gem-miR-15a, that integrates tumor-suppressive miR-15a activity with chemotherapeutic modifications and tumor-targeting capability. This modified miRNA exhibits potent nanomolar activity across diverse EOC models, including PARP inhibitor-resistant cells, without requiring delivery vehicles. Mechanistically, MTX-5-FU-Gem-miR-15a induces replication stress while suppressing G2/M checkpoint regulators (WEE1 and CHK1), resulting in genomic instability and apoptotic cell death. Transcriptomic and protein-level analyses revealed coordinated suppression of resistance-associated and oncogenic signaling pathways, alongside activation of DNA damage coupled with checkpoint abrogation and potential innate immune response. MTX-5-FU-Gem-miR-15a also demonstrates strong synergy with olaparib and robust antitumor efficacy *in vivo*. These findings establish a multimodal miRNA-based therapeutic strategy that targets replication stress and checkpoint dependency to overcome PARPi resistance in ovarian cancer.

## INTRODUCTION

Ovarian cancer (OC) remains the most lethal gynecological malignancy, with an estimated 313,959 new cases and over 200,000 deaths reported globally every year ^1^. Among all histological subtypes, epithelial ovarian cancer (EOC) is the most common and aggressive, accounting for 85-90% of cases and is associated with poor long-term survival ^2^. Standard first-line treatment involves maximal cytoreductive surgery followed by combination chemotherapy, and while initial response rates are high, more than 70% of patients experience relapse ^3,4^. The recurrent tumors often exhibit resistance to chemotherapy, yielding a five-year survival rate below 45% ^5^.

Recent advances in targeted therapies, including poly (ADP-ribose) polymerase (PARP) inhibitors such as olaparib, niraparib, anti-angiogenic agents e.g., bevacizumab, and immune checkpoint inhibitors such as nivolumab, have improved outcomes in select patient populations ^6–9^. However, their overall clinical benefit remains limited by tumor heterogeneity, an immunosuppressive tumor microenvironment, and the emergence of both intrinsic and acquired resistance. For example, PARP inhibitors (PARPi), such as olaparib, have demonstrated significant efficacy against BRCA-mutated or HR-deficient tumors ^10,11^. They exert their antitumor effects by blocking PARP-mediated single-strand break repair and trapping PARP on DNA, which generates replication-associated double-strand breaks that BRCA-mutant or homologous recombination-deficient tumors are unable to repair efficiently, resulting in synthetic lethality ^6^. However, many EOC cases are BRCA wild-type or homologous recombination-proficient and display intrinsic resistance to PARP inhibitors ^12,13^. Moreover, initially responsive tumors develop resistance to PARPi through restoration of homologous recombination via BRCA reversion mutations, replication fork stabilization, or activation of alternative DDR signaling pathways, including the ATR/CHK1/WEE1 checkpoint axis ^14–16^. This reflects a broader challenge in EOC, where highly adaptive and interconnected signaling networks limit the efficacy of single-target strategies and highlight the need for therapeutic strategies capable of simultaneously modulating multiple oncogenic targets and signaling pathways that govern proliferation, stemness, epithelial-to-mesenchymal transition (EMT), and DNA damage tolerance.

microRNAs (miRNAs) are particularly attractive therapeutic candidates in cancer because, unlike single-agent approaches, they can simultaneously regulate multiple genes and modulate diverse signaling pathways ^17,18^. miRNAs are a class of small (∼22 nucleotides), endogenous, non-coding RNAs. They regulate gene expression post-transcriptionally through sequence-specific binding to the 3′ untranslated regions (3′ UTRs) of target mRNAs, leading to mRNA degradation or translational repression ^19^. Because they often bind with incomplete complementarity, miRNAs can suppress multiple targets across functionally related pathways, making them particularly attractive for cancers characterized by pathway redundancy and resistance to therapy ^20,21^. The biological importance of miRNAs was further underscored by the 2024 Nobel Prize, which recognized the discovery of microRNA and its role in post-transcriptional gene regulation. In cancer, epigenetic dysregulation of miRNAs contributes to tumor initiation, progression, and therapeutic resistance. Also, tumor-suppressor miRNAs are frequently downregulated, and restoring their expression has been shown to suppress multiple oncogenic pathways and sensitize tumor cells to chemotherapy. Yet despite this strong biological rationale, no FDA-approved miRNA-based cancer therapeutics exist, largely because clinical translation has been hindered by instability, inefficient delivery, and limited tumor specificity ^22^.

Hsa-miR-15a (miR-15a), encoded at the miR-15a/16-1 cluster on chromosome 13q14, is a well-characterized tumor suppressor frequently deleted or downregulated across malignancies ^23^. In ovarian cancer, deletion of the miR-15a/16-1 locus has been reported in ∼23.9% of cases and is associated with disease progression and poor clinical outcomes ^24,25^. Consistent with this, miR-15a is markedly downregulated in ovarian cancer cell lines and primary tumors ^26^. Restoration of miR-15a expression has been shown to inhibit tumor cell proliferation, migration, and invasion *in vitro* and *in vivo* ^25^. Mechanistically, miR-15a exerts its tumor-suppressive effects by targeting a network of oncogenic drivers involved in cell cycle regulation, apoptosis, stemness, EMT, and DDR signaling.

Restoration of miR-15a can target several key oncogenic drivers, including polycomb complex protein BMI-1 (BMI1), checkpoint kinase 1 (CHK1), WEE1 G2 checkpoint kinase (WEE1), and Yes-associated protein 1 (YAP1) ^27^. Notably, these targets are also frequently overexpressed and functionally relevant in EOC. For instance, overexpression of BCL-2 has been shown to contribute to chemoresistance in ovarian cancer ^28–30^. Elevated nuclear YAP1 expression correlates with poor prognosis in patients with EOC ^31^. BMI1, a marker of cancer stem cells, is frequently upregulated in ovarian malignancies and plays a critical role in promoting proliferation, self-renewal, EMT, and metastasis ^26^. Both CHK1 and WEE1 are key regulators of the G2/M cell cycle checkpoint and are currently being evaluated as therapeutic targets in clinical trials for ovarian cancer ^32–34^. However, targeting these proteins individually has generally yielded limited efficacy, likely due to compensatory signaling and pathway redundancy.

Tumor-suppressor miRNAs such as miR-15a, due to their pleiotropic targeting ability, offer the potential to modulate multiple oncogenic pathways simultaneously ^35^. To enhance its therapeutic potential while addressing the key limitations of nucleic acid therapeutics – namely instability, poor delivery and limited tumor specificity, we engineered a multimodal miRNA-based therapeutic platform, MTX-5-FU-Gem-miR-15a, by substituting uracil (U) and cytidine (C) residues on the guide strand with the chemotherapeutic nucleoside analogs 5-fluorouracil (5-FU) and gemcitabine (Gem), while conjugating methotrexate (MTX) to the passenger strand (Figure 1A). In this integrated design, 5-FU inhibits thymidylate synthase leading to dTMP depletion and DNA damage, gemcitabine incorporates into DNA causing chain termination and replication arrest while also inhibiting ribonucleotide reductase, and MTX inhibits dihydrofolate reductase (DHFR) resulting in reduced tetrahydrofolate pools and impaired nucleotide biosynthesis ^36–38^. Together, these agents disrupt nucleotide metabolism, DNA replication, and cell cycle progression, while preserving miRNA-mediated gene regulation. These modifications are designed to enhance therapeutic potency, enable efficient cellular uptake without lipid-based delivery vehicles, while preserving miRNA-mediated gene regulation ^39^. Importantly, this construct is also expected to release low-doses of 5-FU and gemcitabine, which may retain therapeutic activity while reducing the systemic toxicity typically associated with conventional chemotherapy. Beyond its cytotoxic role, low-dose 5-FU and gemcitabine has also been reported to promote immune activation, further supporting the therapeutic potential of this design ^40,41^. MTX has high affinity for folate receptor alpha (FRα), which is overexpressed in nearly 80% of epithelial ovarian cancer (EOC) patients ^42^. Accordingly, MTX may improve tumor specificity by targeting FRα, enhancing receptor-mediated uptake and reducing off-target effects ^43^. In addition, low-dose MTX has also been reported to induce cGAS-STING pathway, thereby promoting type I interferon signaling and enhancing anti-tumor immune responses, further strengthening the rationale for the design ^44^.

**Figure 1.**
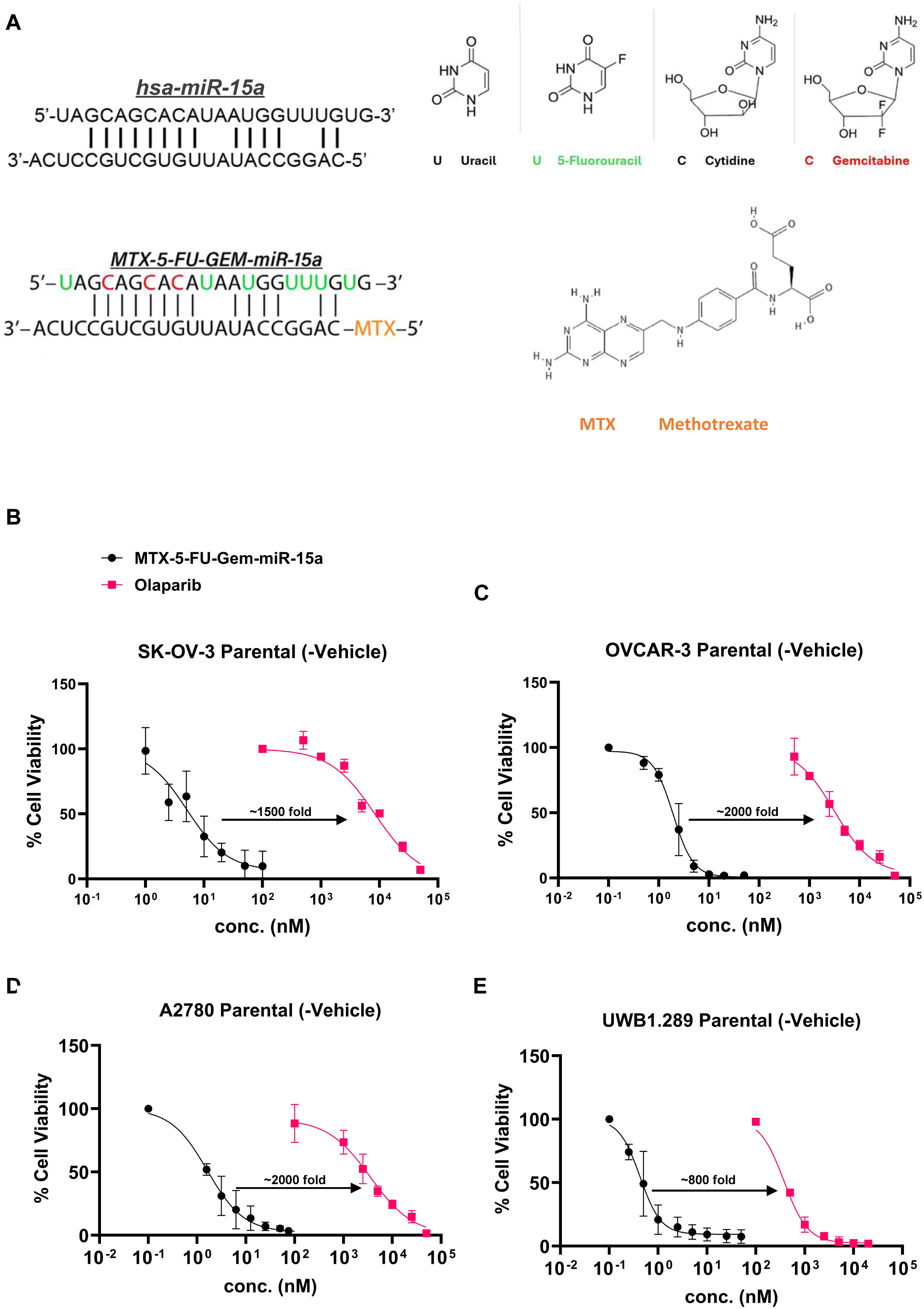
Design of MTX-5-FU-Gem-miR-15a and cytotoxic activity across ovarian cancer models. (A) Schematic representation of hsa-miR-15a and MTX-5-FU-Gem-miR-15a, showing incorporation of 5-fluorouracil and gemcitabine into the miR-15a backbone and conjugation of methotrexate (MTX) to the passenger strand. (B-E) Dose-response curves showing cell viability following treatment with MTX-5-FU-Gem-miR-15a, unmodified miR-15a, and olaparib in SK-OV-3 (B), OVCAR-3 (C), A2780 (D), and UWB1.289 (E) cells. Data represent mean ± SD from n = 4 biological replicates.

In this study, we investigated the efficacy of MTX-5-FU-Gem-miR-15a in both parental and PARPi-resistant epithelial ovarian cancer models. Our results demonstrate that this modified miRNA mimic robustly suppresses cell proliferation without delivery vehicles, induces cell cycle arrest, and promotes apoptotic cell death. Importantly, MTX-5-FU-Gem-miR-15a appears to retain miRNA-mediated target specificity, while the chemotherapeutic molecules contribute to enhanced cytotoxicity and eliminate the need for lipid-based delivery vehicles. Mechanistically, this platform is consistent with replication stress induction coupled with checkpoint disruption and oncogenic pathway suppression, resulting in genomic instability and tumor cell death. Furthermore, MTX-5-FU-Gem-miR-15a exhibits synergistic activity with olaparib and robust antitumor efficacy *in vivo*. These findings support a proof-of-concept for a rationally engineered, multimodal modified miRNA mimic that simultaneously targets oncogenic pathways, disrupts resistance mechanisms, and promotes cell death in tumor cells.

## RESULTS

### MTX-5-FU-Gem-miR-15a achieves nanomolar cytotoxicity across genetically diverse epithelial ovarian cancer cells under vehicle-free condition

To determine the cytotoxic efficacy of MTX-5-FU-Gem-miR-15a in ovarian cancer cells, a dose-response assay was performed across a panel of epithelial ovarian cancer (EOC) cell lines under delivery vehicle-free conditions. The panel included SK-OV-3 (BRCA1/2-WT), OVCAR-3 (BRCA1/2-WT, FRα-high), A2780 (BRCA1/2-WT), and UWB1.289 (BRCA1-null) - spanning HR-proficient and HR-deficient backgrounds and varying TP53 status. Cells were exposed to increasing concentrations of the modified miRNA mimic for 6 days, and cell viability was quantified using the WST-1 assay.

The modified miRNA exhibited potent cytotoxic activity across the entire cell line panel, with sub- to low-nanomolar half-maximal inhibitory concentrations (IC_50_) across all lines – 5.7 nM (SK-OV-3), 1.9 nM (OVCAR-3), 1.7 nM (A2780) and 0.5 nM (UWB1.289) (Figure 1B-E). In contrast, unmodified miR-15a failed to reduce viability in the absence of a delivery vehicle, indicating that chemical modification is required for functional intracellular activity (Figure S1A-C). Furthermore, MTX-5-FU-Gem-miR-15a was approximately 1000 to 2000 fold more potent than the clinically approved PARP inhibitor, Olaparib, which exhibited IC_50_ values in the micromolar range for BRCA-WT cells (∼8.5 µM in SK-OV-3; ∼3.3 µM in OVCAR-3) and a high nanomolar value for the BRCA1-null cells (∼400 nM in UWB1.289).

To further dissect the contribution of the chemotherapeutic components, dose-response analyses were performed using combinations of 5-FU and gemcitabine (7:3) and with the addition of methotrexate (7:3:1). These combinations exhibited cytotoxic effects only at high nanomolar to micromolar concentrations and did not recapitulate the nanomolar potency observed with MTX-5-FU-Gem-miR-15a (Figure S1D-E), indicating that the enhanced efficacy of the construct is not attributable to simple additive effects of the individual drugs.

### MTX-5-FU-Gem-miR-15a induces S-phase accumulation and apoptosis in parental EOC cells

To elucidate the mechanism underlying the cytotoxicity of MTX-5-FU-Gem-miR-15a, we next investigated its effects on cell cycle progression and induction of apoptosis. Following 72 hours of treatment, flow cytometric analysis demonstrated that MTX-5-FU-Gem-miR-15a significantly increased S-phase accumulation with a corresponding reduction in the G₂/S ratio (Figure 2A-B). This effect was substantially greater than that observed with an equimolar concentration of either the unmodified miR-15a mimic or a combination of free 5-FU and gemcitabine. For example, in SK-OV-3 cells, the modified mimic produced a ∼10-fold decrease in the G₂/S ratio (p <0.0001), whereas the free drug combination caused only a 2.5-fold reduction (p = 0.0215). A similar pattern of cell cycle disruption was observed across all EOC cell lines, including OVCAR-3 (∼2.5 fold decrease in G2/S, p = 0.0018), A2780 (∼2.5 fold decrease in G2/S, p <0.0001), and UWB1.289 (∼6 fold decrease in G2/S, p <0.0001), indicating a consistent mechanism of action across diverse genetic backgrounds.

**Figure 2.**
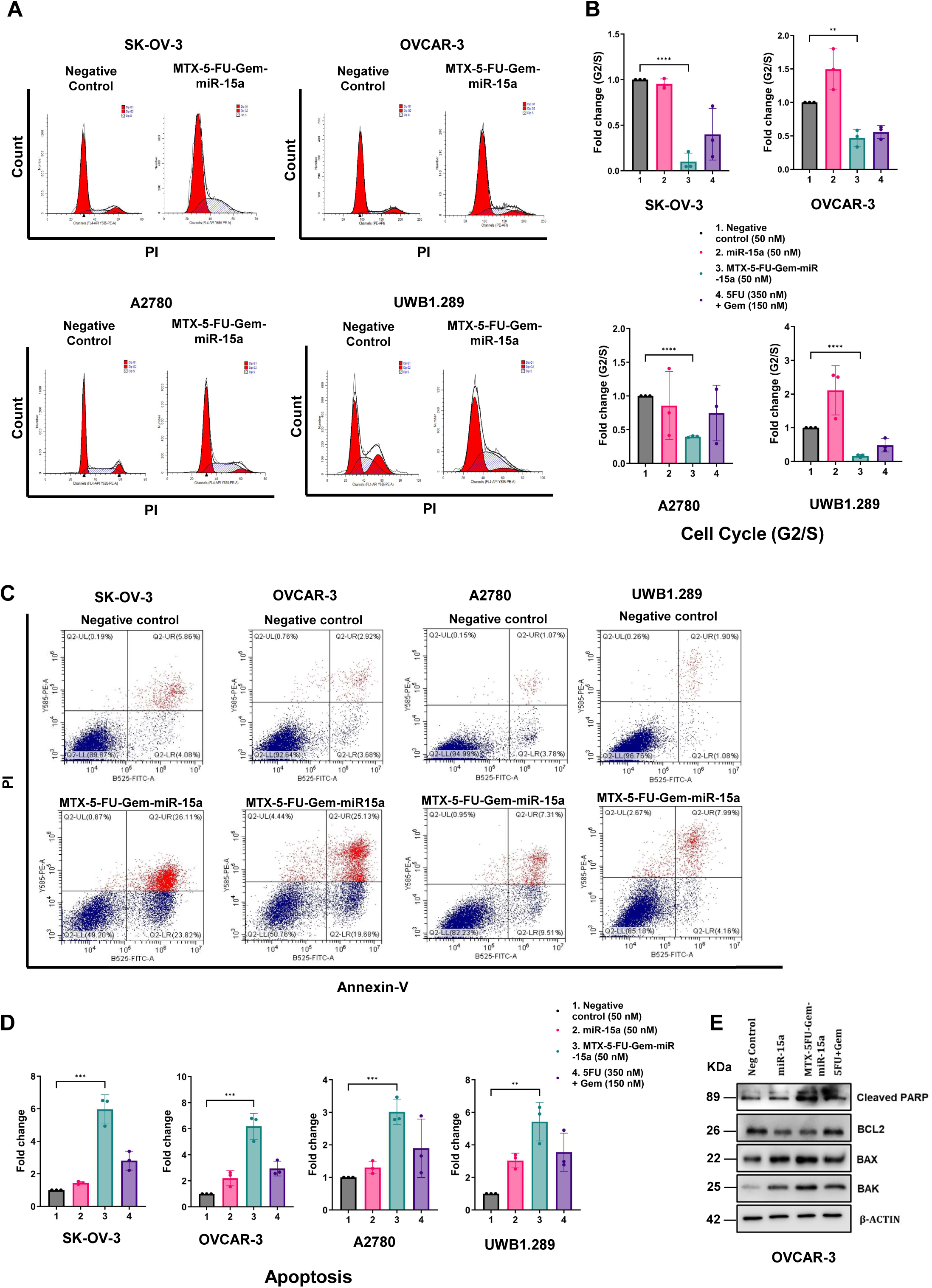
MTX-5-FU-Gem-miR-15a induces S-phase accumulation and apoptosis. (A) Representative histograms of DNA content (PI staining) in only negative control and MTX-5-FU-Gem-miR-15a treatment showing cell cycle distribution in SK-OV-3, OVCAR-3, A2780, and UWB1.289 cells following treatment. (B) Quantification of cell cycle distribution after treatment shown as fold change in G2/S ratio relative to negative control across cell lines. (C) Representative Annexin V/PI flow cytometry plots showing apoptosis in indicated cell lines. (D) Quantification of apoptotic cells (early + late apoptosis) shown as fold change relative to control. (E) Western blot analysis showing expression of cleaved PARP, BCL-2, BAX, and BAK following treatment; β-actin serves as a loading control. Data represent mean ± SD from n = 3 biological replicates. Statistical significance between two groups was determined using two-tailed Student’s t test. ns, not significant; *p < 0.05; **p < 0.01; ***p < 0.001; ****p < 0.0001.

In addition to disrupting cell cycle progression, MTX-5-FU-Gem-miR-15a significantly induced apoptosis in the ovarian cancer cells (Figure 2C-D). Annexin V/PI staining revealed significant increases in total apoptotic population across EOC cell lines - 6-fold in SK-OV-3 (p = 0.0007), 6.2-fold in OVCAR-3 (p = 0.0008), 3-fold in A2780 (p = 0.0009), and 5.5-fold in UWB1.289 (p = 0.003). At the protein level, western blot analysis showed downregulation of the anti-apoptotic protein BCL-2 and a concurrent increase in pro-apoptotic effectors (BAK, BAX, and cleaved PARP) and executioner protein (cleaved caspase-7) (Figure 2E).

These results demonstrate that MTX-5-FU-Gem-miR-15a exerts cytotoxicity through cell cycle disruption and the activation of the apoptotic cell death pathways.

### MTX-5-FU-Gem-miR-15a retains miR-15a target specificity

To confirm that MTX-5-FU-Gem-miR-15a retained its target specificity, we assessed its ability to downregulate established miR-15a targets. Western blot analysis showed that, similar to unmodified miR-15a delivered via transfection, the MTX-5-FU-Gem-miR-15a conjugate markedly reduced the protein expression of key targets, including the G₂/M checkpoint kinases WEE1 and CHK1, which were decreased by approximately 50-55% relative to control (Figure 3A-C). YAP1 and BMI1, regulators of cancer stemness and self-renewal, were similarly downregulated (∼40-50% reduction). The expression of the anti-apoptotic protein BCL-2 and the cell-cycle regulator CCND1 was also significantly decreased by approximately 50-60% compared to control. Quantitative analysis confirmed that the magnitude of target suppression induced by MTX-5-FU-Gem-miR-15a was comparable to that observed with unmodified miR-15a, demonstrating that chemical modification preserves canonical miRNA silencing activity.

**Figure 3.**
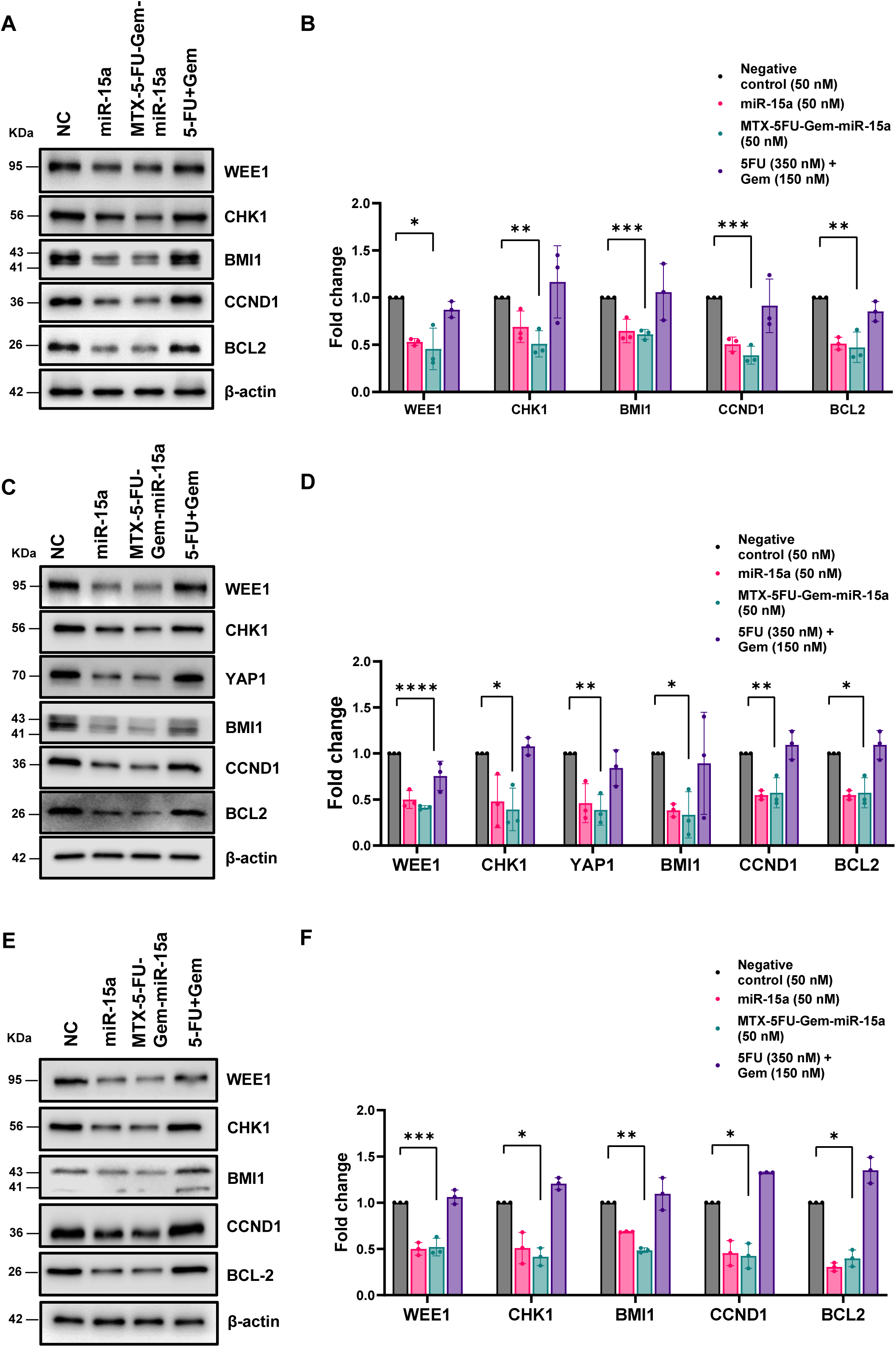
MTX-5-FU-Gem-miR-15a suppresses canonical miR-15a targets. (A, C, E) Western blot analysis of canonical miR-15a target proteins in ovarian cancer cells treated with negative control, miR-15a, MTX-5-FU-Gem-miR-15a, or 5-FU + gemcitabine. Panel A, SK-OV-3; panel C, OVCAR-3; panel E, UWB1.289. (B, D, F) Densitometric quantification of the corresponding western blots normalized to β-actin. Data represent mean ± SD from n = 3 biological replicates. Statistical significance between two groups was determined using two-tailed Student’s t test. ns, not significant; *p < 0.05; **p < 0.01; ***p < 0.001; ****p < 0.0001.

### MTX-5-FU-Gem-miR-15a retains nanomolar potency and induces apoptosis in PARP inhibitor-resistant ovarian cancer cells

To evaluate whether MTX-5-FU-Gem-miR-15a can overcome PARP inhibitor resistance, olaparib-resistant derivatives of SK-OV-3, OVCAR-3, and UWB1.289 ovarian cancer cells were generated by continuous dose-escalation of olaparib (see Methods) and designated SK-OV-3/OlaR, OVCAR-3/OlaR, and UWB1.289/OlaR. The OlaR cells exhibited markedly elevated IC50 values for olaparib exceeding 50 µM in SK-OV-3/OlaR and OVCAR-3/OlaR, and 10 µM in UWB1.289/OlaR, representing ∼2000-fold increases compared to their parental counterparts (Table 1).

Despite this resistance, MTX-5-FU-Gem-miR-15a retained robust cytotoxic activity in all resistant models, with IC50 values of 9.3 nM in SK-OV-3/OlaR, 5.1 nM in OVCAR-3/OlaR, and 1.2 nM in UWB1.289/OlaR, even in the absence of a delivery vehicle (Figure 4A-C). These values are comparable to those observed in parental cells, indicating that olaparib resistance mechanisms do not impair the uptake or activity of the modified miRNA. Furthermore, these data indicate that resistance to PARP inhibition does not confer cross-resistance to the multimodal miRNA therapeutic.

**Figure 4.**
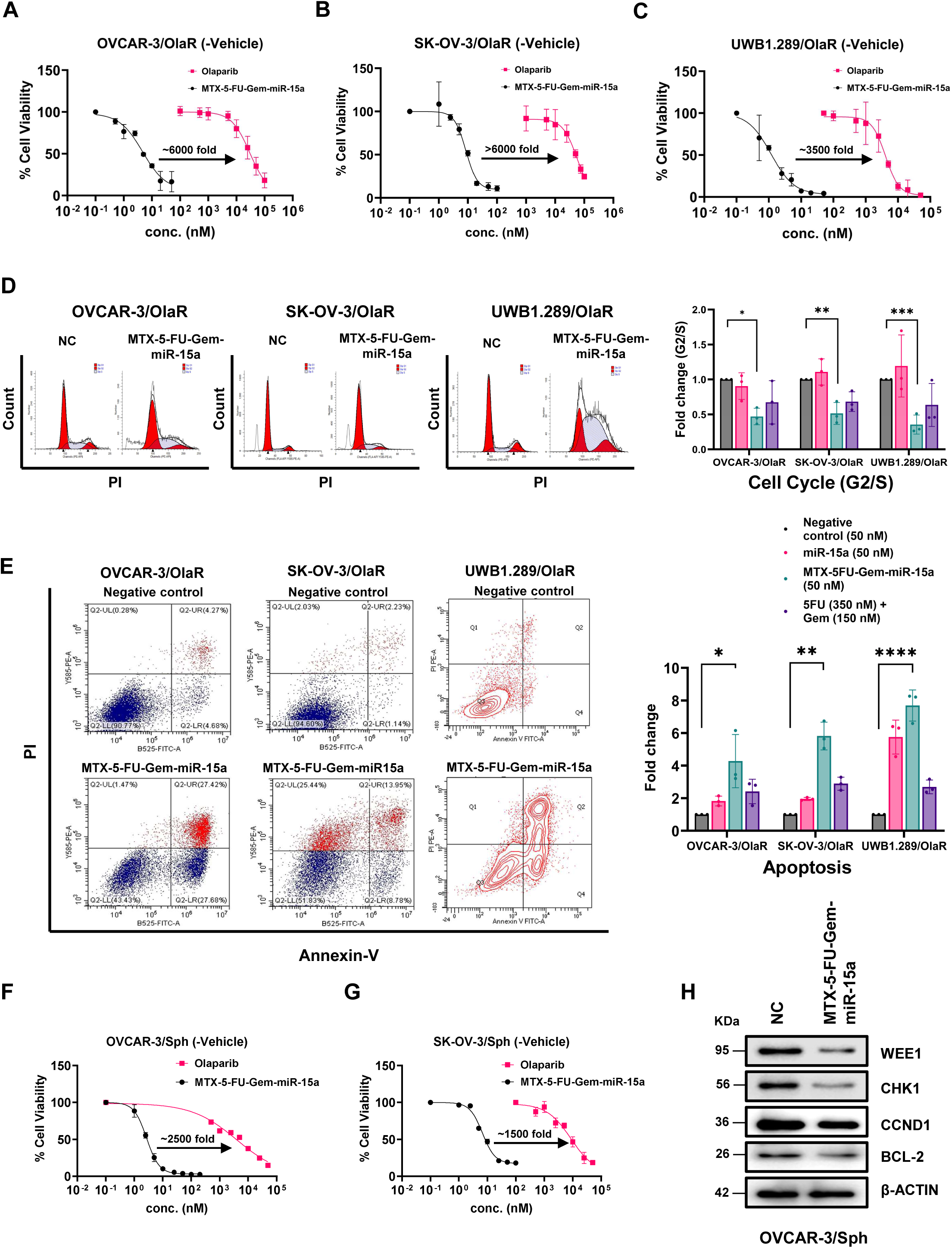
MTX-5-FU-Gem-miR-15a retains activity in PARP inhibitor-resistant models and exhibits effects in 3D cultures. (A-C) Dose-response curves for MTX-5-FU-Gem-miR-15a and olaparib in olaparib-resistant OVCAR-3/OlaR (A), SK-OV-3/OlaR (B), and UWB1.289/OlaR (C) cells. (D) Representative cell cycle histograms (PI staining) and quantification of fold change in G2/S ratio in resistant cell lines. (E) Representative Annexin V/PI plots and quantification of total apoptotic cells (early + late apoptosis). (F-G) Dose-response analysis of cell viability in OVCAR-3 (F) and SK-OV-3 (G) spheroid cultures treated with increasing concentrations of MTX-5-FU-Gem-miR-15a or olaparib. (H) Western blot analysis of WEE1, CHK1, CCND1, and BCL-2 in OVCAR-3 spheroid cultures following treatment; β-actin serves as the loading control. Data represent mean ± SD from n = 3 biological replicates. Statistical significance between two groups was determined using two-tailed Student’s t test. *p < 0.05; **p < 0.01; ***p < 0.001; ****p < 0.0001.

Western blot analysis confirmed that MTX-5-FU-Gem-miR-15a suppressed canonical miR-15a target proteins in olaparib-resistant cells as well, indicating retention of target specificity despite acquired resistance (Figure S2A).

Consistent with parental cells, the modified miRNA mimic treatment produced a significant reduction in the G₂/S ratio across all resistant lines (Figure 4D). Annexin V/PI staining further revealed a substantial increase in apoptotic cell populations across all resistant lines compared to untreated controls (Figure 4E). Together, these findings establish that MTX-5-FU-Gem-miR-15a not only retains its cytotoxic activity in olaparib-resistant ovarian cancer models but also disrupts cell cycle progression and induces apoptosis in a manner consistent with responses observed in parental cells.

### MTX-5-FU-Gem-miR-15a suppresses viability and oncogenic signaling in 3D spheroid models

To assess the therapeutic efficacy in a more physiologically relevant model, we examined MTX-5-FU-Gem-miR-15a in 3D spheroid cultures of OVCAR-3 and SK-OV-3 cells, which are enriched for cancer stem-like cells (CSCs) and known to exhibit drug resistance compared to monolayer culture.

In the absence of any delivery vehicle, MTX-5-FU-Gem-miR-15a potently reduced viability in both OVCAR-3 and SK-OV-3 spheroid cultures, with an IC_50_ of approximately 3 nM and 6 nM - ∼1500 fold lower than olaparib, which required ∼4 µM and ∼10 µM respectively to achieve comparable effects (Figure 4F-G). Representative spheroid images are shown in Figure S2B. Western blot analysis further demonstrated downregulation of WEE1, CHK1, CCND1, and BCL-2 in spheroid cultures treated with MTX-5-FU-Gem-miR-15a (Figure 4H), confirming retention of miR-15a-mediated target suppression in 3D culture.

### MTX-5-FU-Gem-miR-15a induces replication stress and suppresses oncogenic signaling pathways

To elucidate the global transcriptomic changes induced by MTX-5-FU-Gem-miR-15a, RNA sequencing was performed on both parental (PAR) and olaparib-resistant (OR) OVCAR-3 cells following treatment with the MTX-5-FU-Gem-miR-15a (abbreviated M15a in the figures). Subsequent analyses focused on cancer-relevant pathways to better interpret treatment-associated biological effects.

KEGG pathway enrichment analysis revealed positive normalized enrichment score (NES) for DNA damage response and replication stress-associated pathways including Fanconi anemia, homologous recombination, DNA replication, cell cycle, and p53 signaling with concurrent negative NES for metabolic and oncogenic pathways including AMPK, mTOR, PI3K-Akt, and Wnt signaling (Figure 5A). A concordant pattern was observed in olaparib-resistant cells, with positive NES for DNA damage and repair pathways (Fanconi anemia, homologous recombination, DNA replication, p53 signaling, cell cycle) accompanied by negative NES for pro-survival signaling networks, including Wnt, mTOR, PI3K-Akt, and Ras pathways (Figure 5B). Consistent directionality across functionally related pathways in both parental and resistant backgrounds supports the biological relevance of these changes.

**Figure 5.**
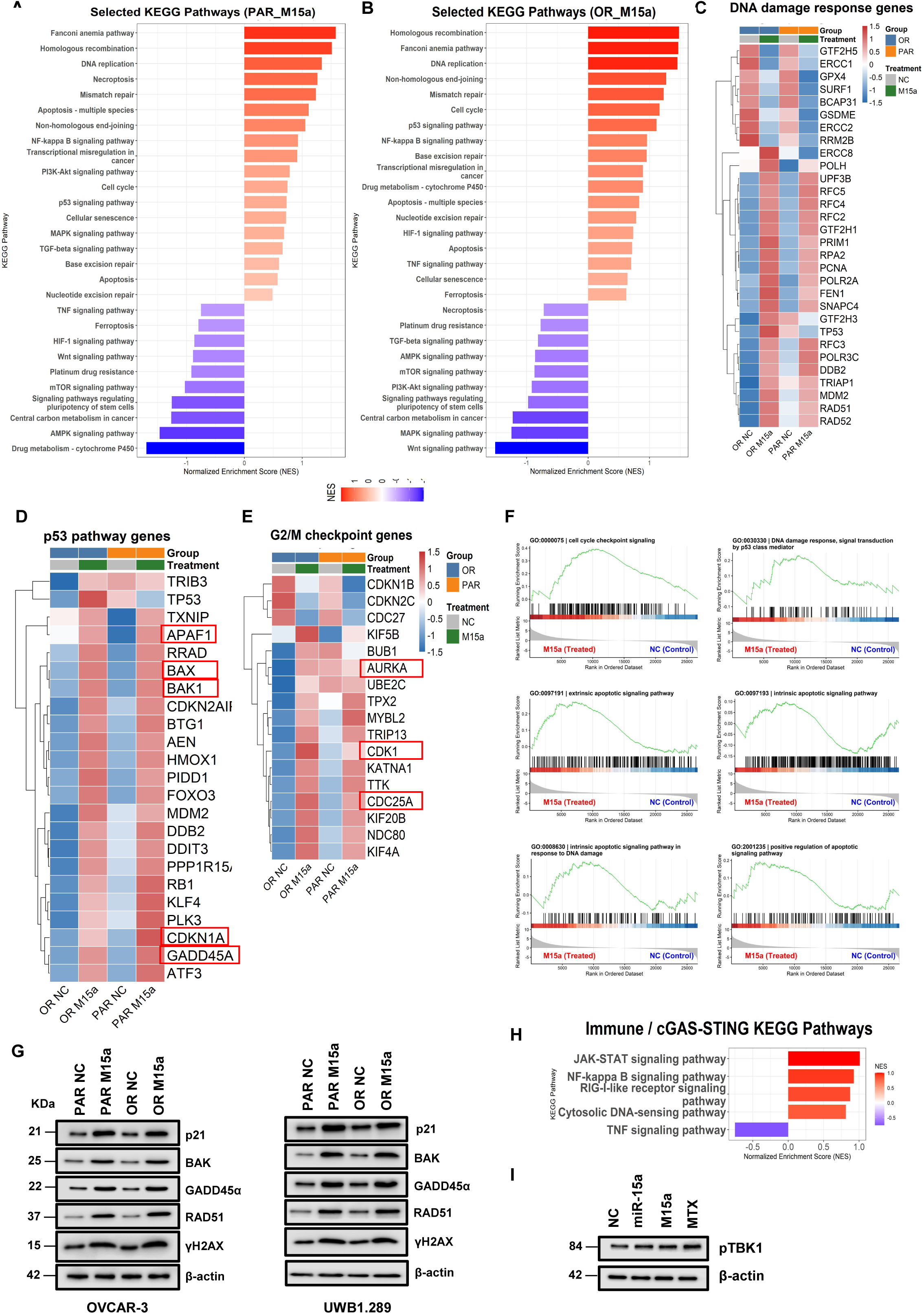
MTX-5-FU-Gem-miR-15a induces replication stress, checkpoint disruption, and immune pathway activation. (A, B) KEGG pathway enrichment analyses in parental (PAR) and olaparib-resistant (OR) OVCAR-3 cells showing enrichment of DNA damage response, cell cycle, and replication-associated pathways, with concurrent suppression of metabolic and oncogenic signaling pathways. (C) Heatmap of DNA damage response and replication-associated genes. (D) Heatmap of p53 pathway genes. (E) Heatmap of G2/M checkpoint genes. (F) Gene set enrichment analysis (GSEA) plots showing enrichment of cell cycle checkpoint signaling, intrinsic apoptotic signaling, apoptotic pathway regulation and p53 signaling pathway in treated versus control conditions. (G) Western blot analysis showing expression of γH2AX, RAD51, GADD45α, BAK, and p21 in parental and resistant cells following treatment; β-actin serves as a loading control. (H) KEGG enrichment analysis of immune-related pathways, including JAK-STAT signaling, NF-κB signaling, RIG-I-like receptor signaling, cytosolic DNA sensing, and TNF signaling. (I) Western blot analysis of pTBK1 following treatment with negative control, miR-15a, M15a, or MTX; β-actin serves as the loading control.

To further characterize pathway-level responses, gene signature scoring using curated Hallmark gene sets was performed across conditions. MTX-5-FU-Gem-miR-15a treatment induced coordinated increases in DNA damage response, replication stress, and p53 pathway activity, along with modulation of G2/M checkpoint-associated gene sets and activation of apoptotic signaling in both parental and resistant cells (Figure S3A). These changes were consistent across both PAR and OR backgrounds, indicating a conserved transcriptional response to treatment.

Hierarchical clustering analysis (heatmaps) revealed upregulation of replication stress-associated genes, including RAD51, RAD52, RPA2, and RFC family members following M15a treatment (Figure 5C). In contrast, several DNA repair and replication maintenance genes such as ERCC1, ERCC2, POLH, PCNA, and RRM2B, were downregulated.

The p53 signaling pathway showed a more nuanced response (Figure 5D). In both PAR and OR cells, MTX-5-FU-Gem-miR-15a treatment led to upregulation of multiple p53-associated genes, including CDKN1A (p21), BAX, and BAK1. Additional stress-responsive and apoptotic regulators, including GADD45A, ATF3, and APAF1, were also increased following treatment.

Analysis of G2/M checkpoint genes demonstrated increased expression of key mitotic regulators, including CDK1, CDC25A, AURKA, and TPX2, in treated conditions relative to controls in both parental and resistant backgrounds, consistent with premature mitotic entry despite unresolved DNA damage (Figure 5E).

Gene set enrichment analysis (GSEA) was further performed to investigate specific, mechanism-relevant pathways, including intrinsic and extrinsic apoptotic signaling, intrinsic apoptotic signaling in response to DNA damage, and p53-mediated signal transduction pathways (Figure 5F). These pathways were selected to provide deeper insight into cell death and stress response mechanisms suggested by earlier analyses. GSEA revealed enrichment patterns in these pathways in treated cells compared to controls, consistent with activation of DNA damage-driven apoptotic signaling.

To examine the relationship between these pathway-level changes, correlation analyses were performed across curated gene sets. Replication stress scores showed a strong positive association with DNA damage signaling, while both replication stress and DNA damage were positively associated with apoptotic pathway activation (Figure S3B-D). Notably, DNA damage signaling also correlated with intrinsic apoptotic pathway activation, consistent with engagement of mitochondrial apoptosis downstream of genomic instability (Figure S3E).

Protein-level validation confirmed activation of DNA damage and apoptotic pathways as found in the transcriptomic analyses (Figure 5G). MTX-5-FU-Gem-miR-15a treatment increased γH2AX and RAD51 levels, along with upregulation of pro-apoptotic markers BAK and p53-associated proteins p21 and GADD45A in both parental and resistant cells, confirming transcriptomic findings at the proetin level.

KEGG pathway analysis further identified enrichment of immune-related signaling pathways, including cytosolic DNA sensing, NF-κB, JAK-STAT, and RIG-I-like receptor following treatment (Figure 5H). Consistent with activation of cGAS-STING signaling pathway, western blot analysis revealed increased phosphorylation of TBK1, supporting activation of the cGAS-STING signaling (Figure 5I).

### MTX-5-FU-Gem-miR-15a reverses PARP inhibitor resistance-associated transcriptional programs and overcomes PARPi resistance

Comparative transcriptomic analysis revealed that MTX-5-FU-Gem-miR-15a treatment reversed resistance-associated gene expression programs in olaparib-resistant EOC cells (Figure 6A-C).

**Figure 6.**
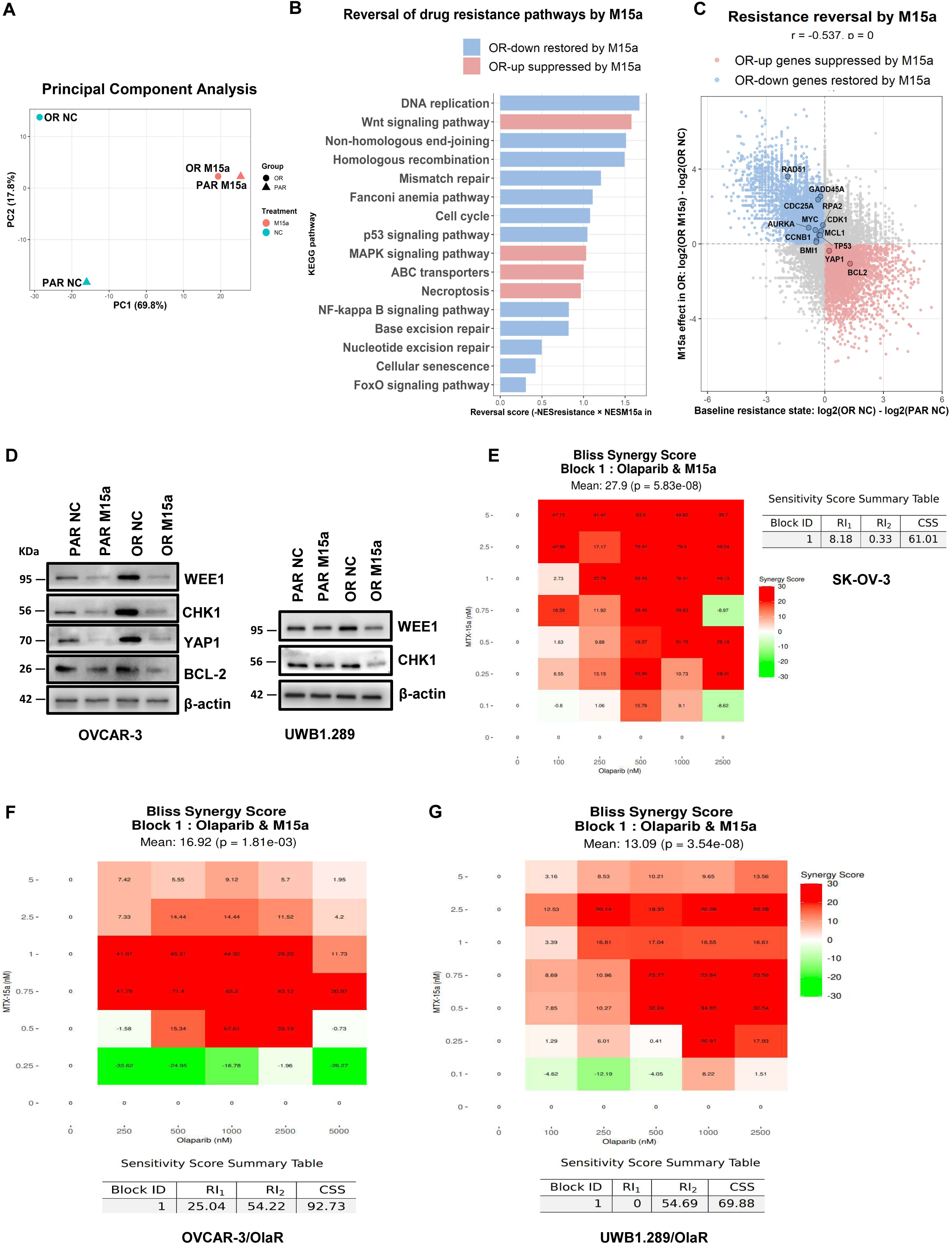
MTX-5-FU-Gem-miR-15a reverses resistance-associated transcriptional and signaling programs. (A) Principal component analysis (PCA) of parental and olaparib-resistant cells under control and treatment conditions. (B) KEGG pathway analysis showing reversal of resistance-associated pathways. (C) Scatter plot showing correlation between baseline resistance-state gene expression and treatment response. (D) Western blot analysis of WEE1, CHK1, YAP1, and BCL-2 in parental and resistant cells after treatment; β-actin serves as loading control. (E-G) Bliss synergy heatmaps and combination sensitivity scores (CSS) for MTX-5-FU-Gem-miR-15a in combination with olaparib.

Principal component analysis demonstrated a clear separation between untreated parental and resistant cells, indicating a global transcriptional shift. However, following treatment, transcriptional profiles of both parental and resistant cells shifted markedly closer together, indicating convergence of transcriptional profiles.

Pathway-reversion analysis identified changes in multiple resistance-associated pathways in a bidirectional pattern (Figure 6B-C). DNA replication, homologous recombination, Fanconi anemia, and cell cycle pathways showed increased reversion scores, indicating broad re-engagement of DNA damage and repair-associated transcriptional programs after modified miRNA treatment. At the same time, pathways elevated in the resistant state were suppressed by treatment, including Wnt signaling, MAPK signaling, ABC transporters – pathways commonly associated with drug efflux, survival signaling, stemness, and therapeutic resistance.

Scatter plot analysis showed a negative correlation between baseline resistance-state gene expression and treatment response (r = -0.537) (Figure 6C). Genes elevated in resistant cells and suppressed after treatment included YAP1, BCL-2, and BMI1, which are associated with resistance, survival, and stemness. Conversely, genes reduced in resistant cells and increased after treatment included DNA damage response and cell cycle related genes such as RAD51, GADD45A, CDC25A, RPA2, CDK1, and AURKA.

Western blot analysis also showed that WEE1 and CHK1 were elevated in olaparib-resistant cells relative to parental controls and were reduced following MTX-5-FU-Gem-miR-15a treatment (Figure 6D), providing protein-level confirmation of transcriptomic changes in key checkpoint regulators.

Collectively, these findings indicate that MTX-5-FU-Gem-miR-15a suppresses resistance-associated survival programs and restores DNA damage-driven stress responses, reprogramming the transcriptional landscape toward a treatment-sensitive state.

### MTX-5-FU-Gem-miR-15a exhibits synergistic activity with olaparib in both intrinsic and acquired resistant ovarian cancer models

Given that MTX-5-FU-Gem-miR-15a remained highly active in resistant models and suppressed checkpoint signaling, we next tested whether it could further sensitize ovarian cancer cells to PARP inhibition.

Using the Bliss independence model, a strong synergistic interaction was observed between MTX-5-FU-Gem-miR-15a and olaparib across all three models, as reflected by mean Bliss synergy scores of 27.9 (SK-OV-3), 16.92 (OVCAR-3/OlaR), and 13.09 (UWB1.289/OlaR) (Figure 6E-G). This indicates robust positive synergy in each system as mean synergy score > 10 indicate synergy. Combination sensitivity scores (CSS) were also high across models, confirming enhanced cytotoxicity relative to either monotherapy.

These data support MTX-5-FU-Gem-miR-15a can potentiate PARP inhibition and may serve as a rational combination partner in both intrinsic and acquired resistance settings.

### MTX-5-FU-Gem-miR-15a suppresses metastatic tumor growth *in vivo* in both parental and resistant models and improves survival

To assess *in vivo* efficacy of MTX-5-FU-Gem-miR-15a, an experimental metastasis model was employed using luciferase-expressing SK-OV-3 and SK-OV-3/OlaR cells in NOD/SCID mice (Figure 7A). Compared to vehicle-treated controls, MTX-5-FU-Gem-miR-15a (3.75 mg/kg, IV, q2d x 8) significantly reduced metastatic tumor burden, as measured by photon flux over time (two-way ANOVA, treatment effect **p < 0.01, treatment x time effect ****p < 0.0001) (Figure 7B-C) and significantly improved overall survival (log-rank p = 0.0494) (Figure 7D). Serum AST and ALT measurements did not reveal significant hepatotoxicity under the treatment conditions used (Figure 7E). In addition, body weight remained stable throughout the treatment period, with no significant treatment-associated loss observed, further supporting favorable tolerability (Figure S4A).

**Figure 7.**
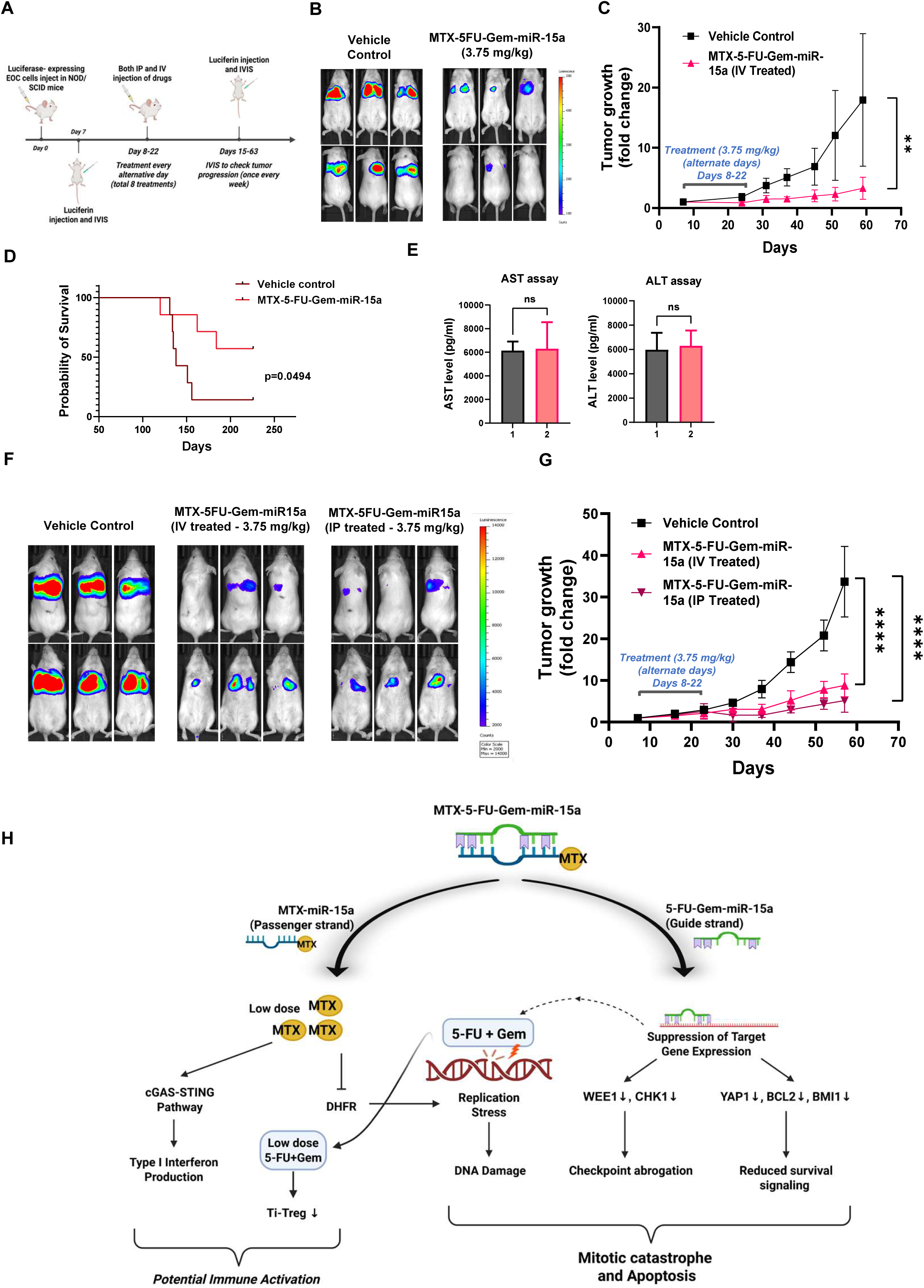
MTX-5-FU-Gem-miR-15a suppresses tumor growth and improves survival *in vivo*. (A) Schematic of the *in vivo* experimental design and treatment schedule. (B) Representative bioluminescence images of parental SK-OV-3 xenografts at endpoint. (C) Tumor growth curves over time for vehicle- and MTX-5-FU-Gem-miR-15a-treated parental xenografts. (D) Kaplan-Meier survival analysis of parental xenografts. (E) Serum AST and ALT measurements evaluating treatment-associated toxicity. (F) Representative bioluminescence images of resistant SK-OV-3/OlaR xenografts treated with vehicle, intravenous MTX-5-FU-Gem-miR-15a, or intraperitoneal MTX-5-FU-Gem-miR-15a. (G) Tumor growth curves over time for resistant xenografts. (H) Proposed mechanistic model of MTX-5-FU-Gem-miR-15a activity. Data represent mean ± SD. Statistical significance for tumor growth and body weight analyses was determined using two-way ANOVA with appropriate post hoc testing. Survival differences were analyzed using the log-rank (Mantel-Cox) test. ns, not significant; *p < 0.05; **p < 0.01; ***p < 0.001; ****p < 0.0001.

In the SK-OV-3/OlaR model, MTX-5-FU-Gem-miR-15a delivered by either the intravenous or intraperitoneal route also robustly suppressed tumor growth relative to vehicle controls (two-way ANOVA; treatment effect ****p < 0.0001, treatment × time effect ****p < 0.0001) (Figure 7F-G), without significant changes in body weight (Figure S4B). Kaplan-Meier analysis did not reveal significant differences in survival between control and treated groups, as the experiment was terminated at a predefined endpoint prior to the occurrence of survival events (Figure S4C).

Together, these results demonstrate that MTX-5-FU-Gem-miR-15a effectively suppresses metastatic tumor growth in both parental and olaparib-resistant EOC models *in vivo*, with evidence of improved survival and no overt systemic toxicity at the dose and schedule tested.

## DISCUSSION

Ovarian cancer remains one of the most lethal gynecologic malignancies, due to its high recurrence rate and the emergence of drug resistance to platinum- or PARP inhibitor-based therapies ^45^. PARP inhibitors, such as olaparib and niraparib, have transformed the management of ovarian cancer, yet both intrinsic and acquired resistance remain pervasive, arising through restoration of homologous recombination, replication fork stabilization, and heightened dependence on ATR/CHK1/WEE1-mediated checkpoint signaling ^10,14–16^. Because the mechanisms are diverse and often co-occur, targeting a single oncogenic pathway has generally provided limited and often transient clinical benefit. Together, these limitations highlight the urgent need for novel therapeutics that can simultaneously induce DNA damage, suppress repair capacity, and promote tumor cell death.

To address this challenge, we developed a multimodal, chemically engineered miRNA mimic, MTX-5-FU-Gem-miR-15a, that integrates the tumor-suppressive activity of miR-15a with the cytotoxic and targeting potential of methotrexate (MTX), 5-fluorouracil (5-FU), and gemcitabine (Gem). The design builds on our previously established platform of nucleoside analog-modified miRNAs, including 5-FU-miR-15a in colon cancer, Gem-miR-15a and Gem-miR-194 in pancreatic cancer, and 5-FU-miR-129 in small cell lung cancer. These studies collectively demonstrated that both 5-FU and gemcitabine modifications enhance miRNA stability and vehicle-free delivery *in vitro*, while conferring additional cytotoxicity ^27,39,46^.

In MTX-5-FU-Gem-miR-15a, 5-FU and gemcitabine modifications on the guide strand preserve miR-15a’s sequence specificity, while MTX conjugation on the passenger strand introduces a folate receptor α (FRα)-targeting module to facilitate tumor-selective uptake. Importantly, these guide-strand modifications do not disrupt Watson-Crick base pairing, thereby preserving target recognition and maintaining miR-15a target specificity without altering its fundamental silencing properties ^47^. Although MTX, 5-FU, and Gem are not standard treatments for ovarian cancer, their incorporation was driven by mechanistic and chemical considerations. These are nucleoside analog chemotherapeutics that target fundamental processes of nucleotide metabolism and DNA synthesis and can be chemically integrated into the guide strand without altering sequence specificity or target recognition^39,48^. Following cellular internalization, the conjugate is expected to undergo degradation by both 5′ and 3′ exonucleases, releasing little but biologically active amounts of 5-FU, Gem, and MTX, thereby providing an additional layer of therapeutic activity. Importantly, this low-dose drug release also offers added therapeutic benefit by reducing systemic toxicity while maintaining anticancer activity and potentially promoting immunostimulatory effects^40,44,49^.

MTX-5-FU-Gem-miR-15a exhibited potent cytotoxic activity across a genetically diverse panel of EOC cell lines, including SK-OV-3 (BRCA-WT, TP53-null, HRP), OVCAR-3 (BRCA-WT, TP53-mutant, HRD, FRα-high), A2780 (BRCA-WT, TP53 WT), and UWB1.289 (BRCA1-null, TP53-null, HRD). This panel of cell lines spans HR-proficient and HR-deficient backgrounds, varying BRCA1/2 mutation status, and both wild-type and mutant TP53. MTX-5-FU-Gem-miR-15a consistently achieved sub- to low-nanomolar IC_50_ across this panel, reflecting potency that substantially exceeds conventional chemotherapeutic agents, which typically require micromolar concentrations to achieve comparable effects (Figure 1B-E; Table 1). This level of potency may reflect the combined contribution of miRNA-mediated target suppression and the incorporated chemotherapeutic modifications, enabling simultaneous engagement of multiple oncogenic and stress response pathways. Notably, although enhanced activity in FRα-overexpressing OVCAR-3 cells is consistent with a role for receptor-mediated uptake due to MTX conjugation, the comparable nanomolar potency observed in other cell lines (SK-OV-3, A2780, and UWB1.289) indicates that FRα-mediated endocytosis is not the sole mechanism of cellular entry. The nucleoside analog modifications themselves facilitate uptake without any delivery vehicle.

Mechanistically, MTX-5-FU-Gem-miR-15a induces a convergence of replication stress and checkpoint disruption that drives tumor cell death. The combined effects of Gem, 5-FU, and MTX disrupt nucleotide metabolism and DNA synthesis, leading to stalled replication forks and activation of the DNA damage response ^37,50^. Gemcitabine-mediated chain termination and depletion of deoxynucleotide pools, together with 5-FU-dependent thymidylate synthase inhibition and methotrexate-mediated blockade of folate metabolism, collectively impair DNA replication and repair ^36–38^. Transcriptomic profiling revealed that MTX-5-FU-Gem-miR-15a treatment induced genes associated with replication stress and homologous recombination, including RAD51, RAD52, RPA2, and components of the Fanconi anemia pathway (Figure 5A-C). Importantly, these transcriptional changes, together with increased γH2AX and RAD51 (Figure 5G) and a pronounced reduction in the G₂/S ratio observed across both parental and resistant cell lines, support a model of S-phase accumulation driven by replication stress. At the same time, the miRNA component downregulates key G2/M checkpoint regulators WEE1 and CHK1 as demonstrated in Figure 6D, which normally delay mitotic entry following DNA damage to provide time to repair the damaged DNA ^51^. By suppressing these gatekeeper kinases, MTX-5-FU-Gem-miR-15a compromises the G2/M checkpoint function and promotes inappropriate cell-cycle progression in the presence of genomic damage. This forced checkpoint bypass is reinforced by RNA-seq evidence of increased CDK1, CDC25A, AURKA, and TPX2 expression. The resulting imbalance between DNA damage burden and checkpoint control is consistent with the pronounced S-phase accumulation, reduced G2/M populations, and robust apoptotic responses observed across both parental and olaparib-resistant ovarian cancer models after modified miRNA treatment (Figures 2 and 4D-E). Together, these findings support a model in which the chemotherapeutic components generate a level of replication damage that exceeds cellular repair capacity, while the miRNA component removes the checkpoint safeguards that would normally allow cells to pause and repair – a combination that ultimately results in mitotic catastrophe and cell death, as summarized in our proposed mechanistic model (Figure 7H) ^52,53^.

This dual mechanism directly addresses the clinical challenge of resistance to PARP inhibitor therapy. PARP inhibitor-resistant ovarian cancers typically retain high levels of replication stress but survive through a combination of replication fork stabilization, restoration of homologous recombination repair, and increased dependence on ATR-CHK1-WEE1 – mediated cell cycle checkpoints ^16,45,54,55^. This checkpoint dependency represents a conserved vulnerability in both intrinsically resistant HRP tumors and HRD tumors that acquire resistance through restoration of DNA repair capacity ^56,57^. Our findings indicate that MTX-5-FU-Gem-miR-15a exploits this vulnerability to overcome these resistance mechanisms retaining nanomolar IC₅₀ values across all resistant lines (Figure 4A-C; Table 1). Transcriptomic analyses showed that treatment partially reprograms the resistant transcriptome, with convergence of parental and resistant profiles and suppression of pathways linked to survival and drug resistance, including Wnt, MAPK, and ABC transporter signaling, as revealed by pathway-reversion analysis (Figure 6B). These pathways are well established drivers of therapeutic resistance, where Wnt signaling contributes to cancer stemness and survival, and ABC transporters mediate drug efflux and reduced intracellular drug accumulation ^58–60^. Their downregulation following treatment is therefore consistent with reduced resistance capacity and may enhance intracellular retention of therapeutic agents, potentially sensitizing tumor cells to both the modified miRNA and co-administered therapies such as PARP inhibitors. At the gene level, reversal of resistance-associated regulators such as BCL-2 and YAP1 further supports this reprogramming, as these factors are known to promote survival, stemness, and drug resistance in ovarian cancer ^28,30,61,62^. Similarly, restoration of mitotic driver gene expressions such as CDK1, CDC25, AURKA etc. is consistent with disruption of checkpoint-dependent survival states, reinforcing the proposed model of checkpoint abrogation. Western blot analysis further supported this as WEE1 and CHK1, which were elevated in resistant cells, were reduced following treatment (Figure 6D). This phenotypic outcome is conceptually similar to induced synthetic lethality observed with pharmacological ATR or WEE1 inhibitors in preclinical ovarian cancer models but is achieved here through a single multifunctional therapeutic.

Consistent with this mechanism, MTX-5-FU-Gem-miR-15a exhibited strong synergistic activity when combined with olaparib in EOC cells with intrinsic or acquired resistance to PARPi, with Bliss synergy scores indicating robust positive interaction across all models tested (Figure 6E-G). This synergy is mechanistically coherent, as olaparib-mediated PARP trapping increases reliance on replication stress response pathways, while MTX-5-FU-Gem-miR-15a simultaneously amplifies replication stress and suppresses checkpoint signaling ^56^. Together, these effects create an unsustainable level of genomic instability, forcing cells to progress through the cell cycle with unresolved DNA damage and ultimately triggering mitotic catastrophe. These findings position MTX-5-FU-Gem-miR-15a not only as a monotherapy for PARP inhibitor-resistant disease, but also as a rational combination partner to enhance and prolong the efficacy of PARP inhibitors in ovarian cancer.

A further dimension of the therapeutic mechanism is its independence from TP53 status. The extensive genotoxic stress is associated with activation of a p53-associated transcriptional program, characterized by increased expression of CDKN1A (p21), GADD45A, and the pro-apoptotic effectors BAX, BAK1, and APAF1 (Figure 5D). Strikingly, these molecular signatures were observed in TP53-mutant OVCAR-3 cells, indicating that MTX-5-FU-Gem-miR-15a can trigger apoptotic signaling independent of functional p53. The induction of p21 and mitochondrial apoptotic mediators in modified miRNA-treated cells suggests that alternative stress-responsive transcription factors and checkpoint regulators may be sufficient to drive cell-cycle arrest and programmed cell death in the absence of canonical p53 activity ^63,64^. Collectively, these findings suggest that MTX-5-FU-Gem-miR-15a can induce irreversible DNA damage and apoptosis regardless of TP53 status, a key advantage in the treatment of high-grade serous ovarian cancers (HGSOC) where TP53 mutations are nearly universal ^65,66^.

Beyond its effects on DNA damage and cell cycle regulation, MTX-5-FU-Gem-miR-15a was also observed to suppress major oncogenic and metabolic pathways implicated in ovarian cancer progression and therapeutic resistance. Pathway enrichment analyses revealed downregulation of PI3K-AKT, mTOR, AMPK, and Wnt signaling which promote proliferation, stemness, and survival ^27,58,67,68^. Key regulators in these pathways, such as BMI1 and YAP1, are validated miR-15a targets and play central roles in maintaining the cancer stem cell (CSC) population ^27^. The simultaneous repression of these pathways may contribute to pronounced reduction in proliferation in both monolayer and 3D spheroid cultures. Importantly, retention of nanomolar potency in 3D spheroids is consistent with activity in more therapy-resistant, stem-like cellular states that contribute to recurrence (Figure 4F-G) ^69,70^.

An additional and particularly promising feature of MTX-5-FU-Gem-miR-15a lies in its potential to modulate antitumor immunity through localized release of low levels of methotrexate and gemcitabine. Prior studies have shown that low-dose methotrexate can activate the cGAS-STING pathway and enhance type I interferon production^44^. Consistent with this, RNA-seq and protein analyses demonstrate enrichment of innate immune-related pathways and increased pTBK1 levels are consistent with cGAS-STING signaling (Figure 5H-I). Similarly, low-dose 5-FU and gemcitabine has been shown to promote immunogenic cell death, enhance CD8⁺ T-cell and NK-cell activity, and reduce myeloid-derived suppressor cells (MDSCs) ^40,41^. In addition, low-dose Gem can deplete the tumor-infiltrating regulatory T cell (TI-Tregs) populations, potentially shifting the tumor microenvironment toward a more immunostimulatory state ^71^. Although these effects were not functionally evaluated in the present study, they raise the possibility that MTX-5-FU-Gem-miR-15a could exert immunomodulatory effects *in vivo* and may synergize with immune checkpoint blockade.

Finally, the therapeutic potential of MTX-5-FU-Gem-miR-15a was confirmed *in vivo*, where treatment at 3.75 mg/kg significantly suppressed metastatic tumor burden in both parental and olaparib-resistant SK-OV-3 models and improved survival in the parental model as shown in Figure 7. Notably, this dosing is substantially lower than the 30-50 mg/kg range commonly used for conventional chemotherapeutic agents in mouse models, and no significant hepatotoxicity or body weight loss was observed, supporting a favorable therapeutic index. Importantly, this favorable safety profile is also consistent with prior clinical study using a related 5-FU modified miR-15a therapeutic, which was well tolerated in patients with relapsed or refractory AML and demonstrated target modulation and disease stabilization, with initial signs of clinical efficacy ^72^. This reduced dosing requirement, together with the observed nanomolar potency *in vitro*, supports the potential for an improved therapeutic index, although further pharmacokinetic and toxicity studies will be required to fully evaluate this.

Despite these promising results, several limitations of the current study remain to be addressed. While methotrexate conjugation was designed to target FRα-overexpression in ovarian cancer, direct validation of FRα-dependent tumor selectivity is required ^43^. Future studies should also characterize the pharmacokinetics, tumor selectivity, and biodistribution of the modified miRNA. In addition, the proposed immunomodulatory effects suggested by transcriptomic and pTBK1 data remain speculative and should be evaluated in immunocompetent mouse models. Finally, although we demonstrated robust efficacy in cell line-based and *in vivo* metastatic models, validation in patient-derived organoids and xenografts across diverse molecular subtypes of EOC will be important to further establish translational relevance.

In conclusion, MTX-5-FU-Gem-miR-15a represents a new platform of multimodal miRNA-based therapeutics that integrates RNA interference with chemotherapeutic activity and tumor-targeting capability. Through coordinated induction of replication stress, abrogation of G2/M checkpoint control, activation of apoptotic signaling, and suppression of resistance-associated transcriptional programs, inhibition of oncogenic survival pathways, MTX-5-FU-Gem-miR-15a has the potential to reprogram PARPi-resistant EOC cells and drive tumor cell death, with additional potential to modulate anti-tumor immunity. Given its activity across both BRCA-WT and BRCA-mutant contexts, these findings provide a framework for the clinical development of multimodal miRNA therapeutics as both monotherapy and rational combination partner with PARP inhibitors in epithelial ovarian cancer.

## RESOURCE AVAILABILITY

### Lead contact

Requests for further information and resources should be directed to and will be fulfilled by the lead contact, Dr. Jingfang Ju.

### Materials availability

MTX-5-FU-Gem-miR-15a and related reagents generated in this study will be made available by the lead contact upon reasonable request and may require a completed materials transfer agreement.

### Data and code availability

- RNA sequencing data generated in this study will be deposited in GEO and will be publicly available as of the date of publication.
- All other data supporting the findings of this study are available from the lead contact upon request.
- All original code used for RNA sequencing and downstream analyses will be made publicly available upon publication.
- Any additional information required to reanalyze the data reported in this paper is available from the lead contact upon request.

## ACKNOWLEDGMENTS

This work was supported by the VA Merit Award BX005260-01 (J. Ju). We thank members of the Ju laboratory and the Department of Pathology for helpful discussions and technical assistance.

## AUTHOR CONTRIBUTIONS

Conceptualization, A.P. and J.J.; methodology, A.P., A.O., H.B., L.C., and I.O.; investigation, A.P. and A.O.; formal analysis (RNA sequencing and bioinformatics analysis), A.P.; writing—original draft, A.P.; writing—review and editing, A.P., A.O., and J.J.; supervision, J.J.; funding acquisition, J.J.; resources, J.J.

## DECLARATION OF GENERATIVE AI AND AI-ASSISTED TECHNOLOGIES

During the preparation of this work, the authors used ChatGPT (OpenAI) to assist with language editing. All content was reviewed and edited by the authors, who take full responsibility for the final manuscript.

## SUPPLEMENTAL INFORMATION

**Document S1.** Figures S1-S4, and Table S1

**Table S2.** Differential gene expression analysis of parental and olaparib-resistant OVCAR-3 cells following MTX-5-FU-Gem-miR-15a treatment, related to Figure 5 and Figure 6.

## TABLES AND TEXT BOXES

IC_50_ values of MTX-5-FU-Gem-miR-15a and olaparib across parental, olaparib-resistant, and spheroid ovarian cancer models.

| Cell line | MTX-5-FU-Gem-miR-15a<br>IC<sub>50</sub> (nM) | Olaparib<br>IC<sub>50</sub> (nM) | Fold change |
| --- | --- | --- | --- |
| OVCAR-3 | 1.89 ± 0.31 | 3288 ± 2000 | ~ 2000 |
| SK-OV-3 | 5.71 ± 4.86 | 8651 ± 4300 | ~ 1500 |
| A2780 | 1.69 ± 0.59 | 2972 ± 1650 | ~ 2000 |
| UWB1.289 | 0.47 ± 0.12 | 386.75 ± 132.9 | ~ 800 |
| OVCAR-3/OlaR | 5.14 ± 2.21 | ~ 30000 | ~ 6000 |
| SK-OV-3/OlaR | 9.34 ± 1.7 | > 50000 | > 6000 |
| UWB1.289/OlaR | 1.22 ± 0.52 | 3989 ± 1142 | ~ 3500 |
| OVCAR-3/Sph | 2.8 ± 0.22 | 3989 ± 1140 | ~ 2500 |
| SK-OV-3/Sph | 6.7 ± 0.5 | 10250 ± 3300 | ~ 1500 |

## STAR★METHODS

## KEY RESOURCES TABLE

See the more detailed <u>Word table template</u> document for examples of how to list items.

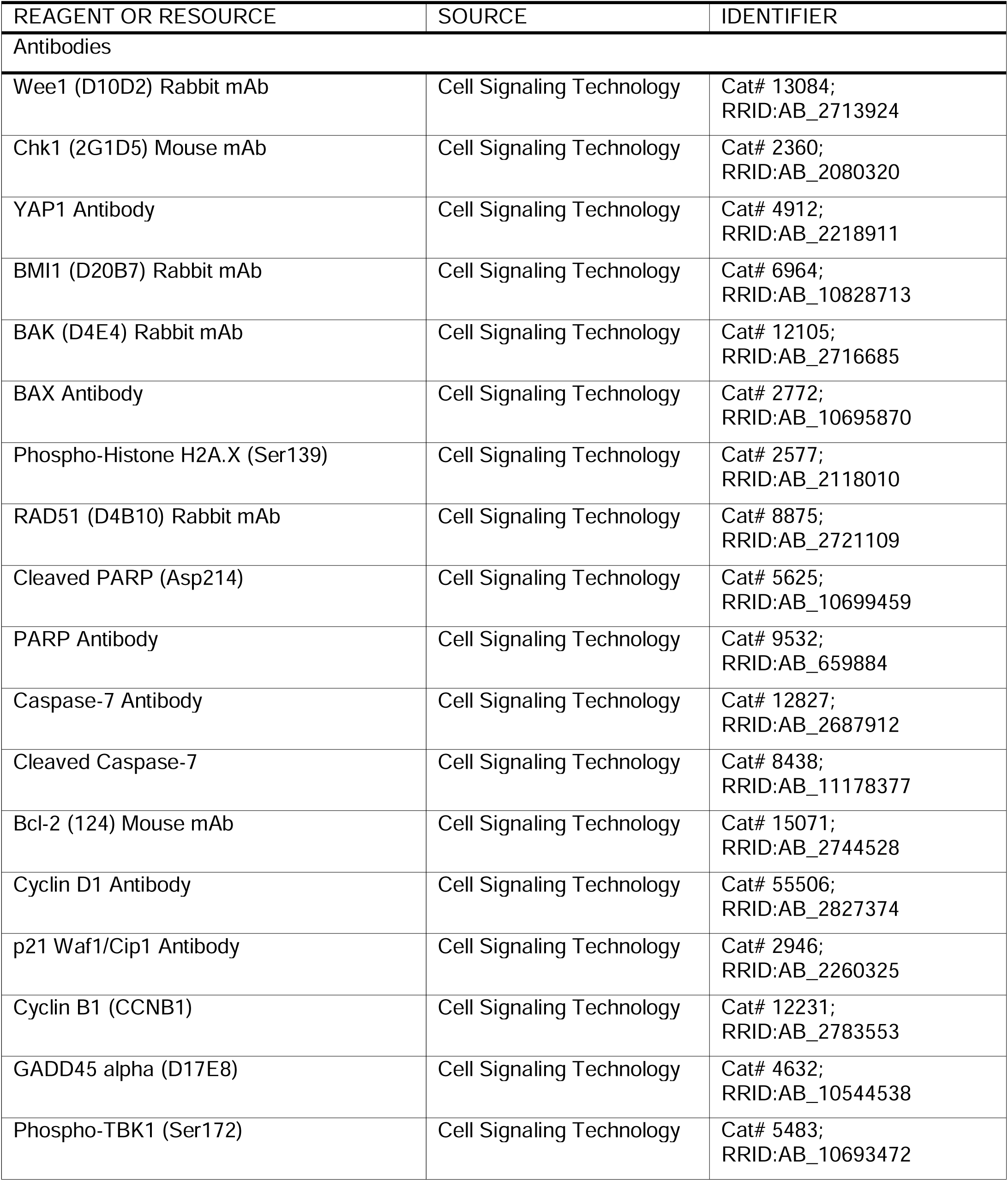
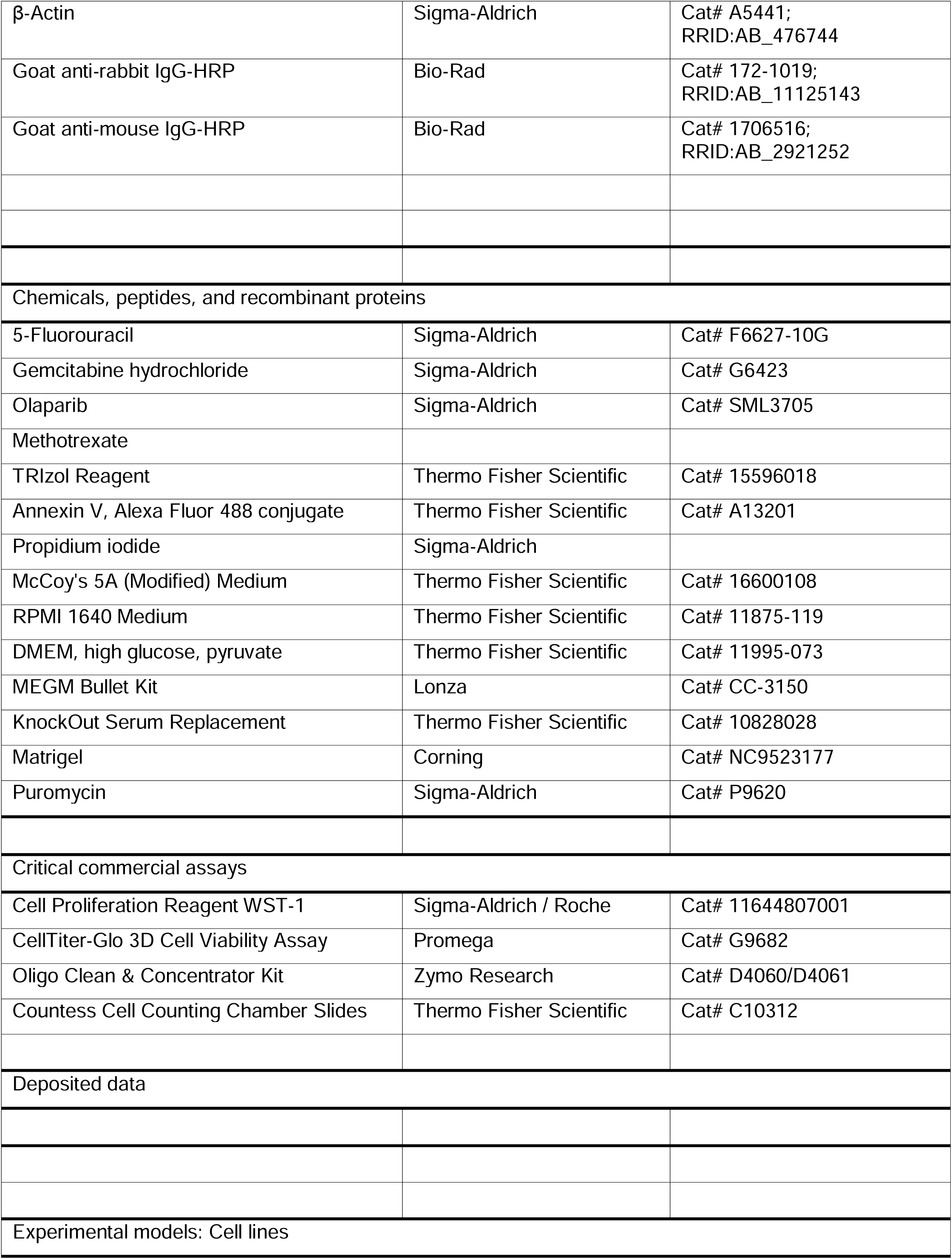
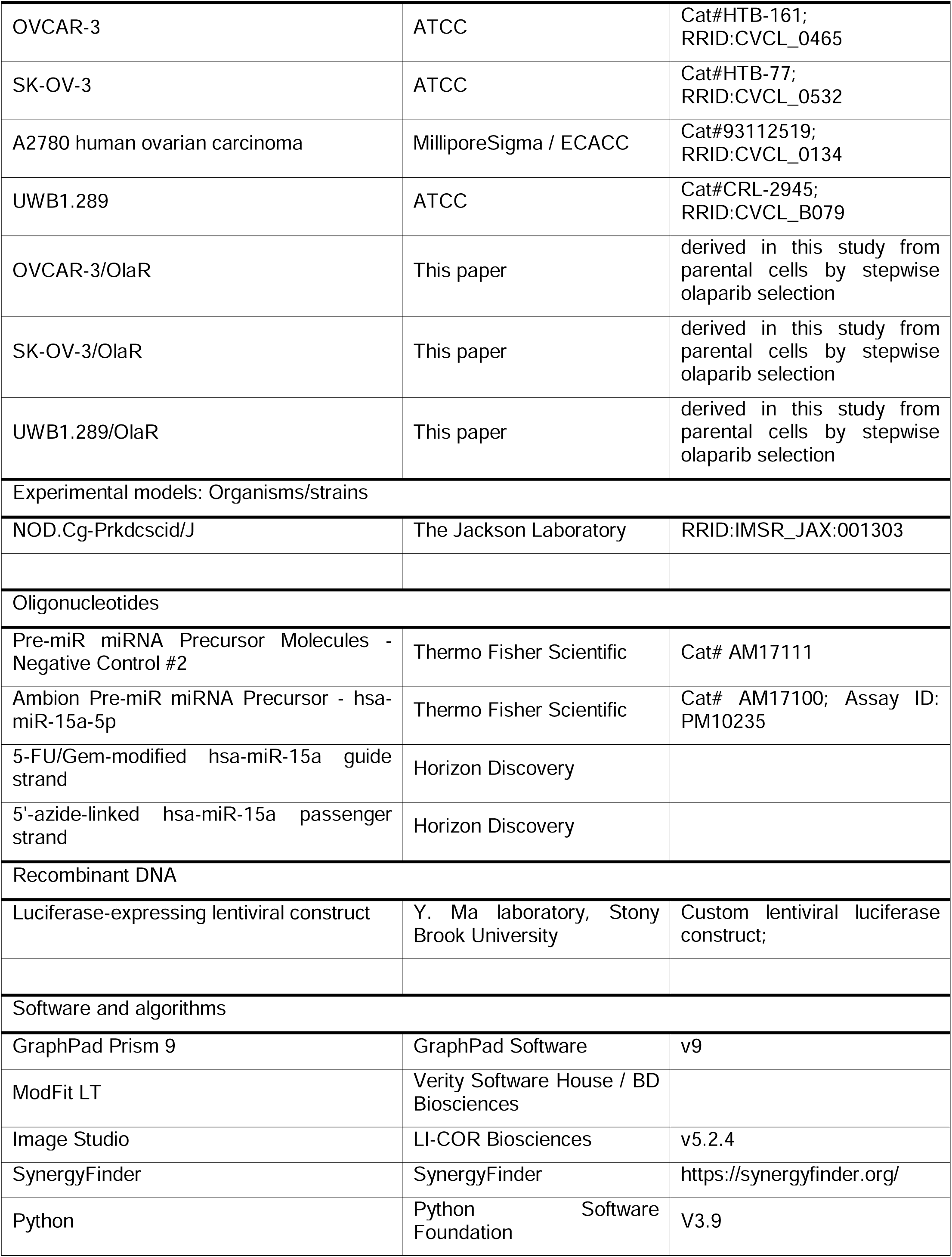
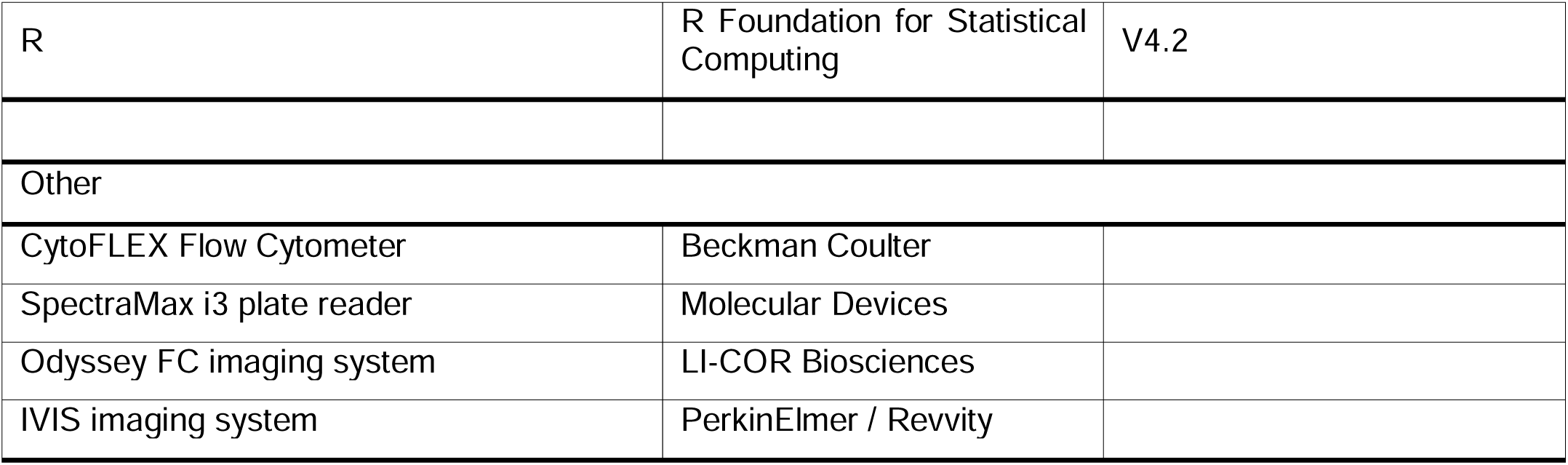

## EXPERIMENTAL MODEL AND STUDY PARTICIPANT DETAILS

### Cell Lines and Culture Conditions

Human epithelial ovarian cancer (EOC) cell lines SK-OV-3, OVCAR-3, A2780, and UWB1.289 were obtained from the American Type Culture Collection (ATCC) and MilliporeSigma. Cells were maintained at 37°C in a humidified incubator with 5% CO₂. SK-OV-3 cells were cultured in McCoy’s 5A medium supplemented with 10% fetal bovine serum (FBS), OVCAR-3 cells in RPMI-1640 supplemented with 10% FBS and 0.01 mg/mL bovine insulin, A2780 cells in RPMI-1640 supplemented with 10% FBS and 2 mM L-glutamine, and UWB1.289 cells in RPMI-1640 supplemented with 3% FBS and 50% Mammary Epithelial Cell Growth Medium (MEGM) BulletKit. Prior to addition, FBS was sterilized via Steriflip-GP, 0.22 µm (MilliporeSigma, St. Louis, MO, USA). Cell line identities were confirmed by short tandem repeat profiling, and Mycoplasma testing was negative.

### Generation of Olaparib-Resistant Cell Lines

For the development of olaparib-resistant models, OVCAR-3, SK-OV-3, and UWB1.289 cells were chosen to address clinically relevant ovarian cancer subtypes with distinct DNA repair capacities. Specifically, OVCAR-3 (BRCA1-WT, HR-deficient), SK-OV-3 (BRCA -WT, HR proficient, multidrug resistant), and UWB1.289 (BRCA1-null, HR-deficient) provide complementary models for studying acquired PARP inhibitor resistance. To make them olaparib-resistant, the cell lines were exposed to gradually increasing concentrations of olaparib (MilliporeSigma, St. Louis, MO, USA). OVCAR-3 and SK-OV-3 cells were sequentially treated with 10-100 µM olaparib, whereas UWB1.289 cells were exposed to 0.5-10 µM olaparib. Drug concentrations were increased in a stepwise manner once cells regained normal proliferation rates at the prior concentration. This selection process was carried out over several months until stable resistant populations were established, hereafter referred to as OVCAR-3/OlaR, SK-OV-3/OlaR, and UWB1.289/OlaR. Resistant lines were maintained in drug-free medium for at least two passages before use in experiments to minimize acute drug-response effects.

### Luciferase-Expressing Cell Lines

The luciferase-expressing EOC cell lines, SK-OV-3-Luc and SK-OV-3-Luc/OlaR, were generated by transfecting SK-OV-3 and SK-OV-3/OlaR cells with a luciferase-expressing lentiviral construct (obtained from the Y. Ma laboratory at the Renaissance School of Medicine at Stony Brook University). Following successful transfection, luciferase-positive cells were selected using puromycin in McCoy’s 5A (Modified) medium (Thermo Fisher Scientific, Waltham, MA, USA). After stable selection, the cell lines were maintained in McCoy’s 5A (Modified) Medium supplemented with 10% fetal bovine serum (FBS) (MilliporeSigma, St. Louis, MO, USA). Luciferase expression was confirmed using an *In vivo* Imaging System (IVIS) prior to use *in vivo* experiments.

### Mice

6-7 weeks old female NOD/SCID mice (JAX: 001303) were purchased from Jackson Laboratory. All the experiments with mice were conducted in Stony Brook University animal care facility and in accordance with the Institutional Animal Care and Use Committee (IACUC).

## METHOD DETAILS

### Design and Synthesis of MTX-5-FU-Gem-miR-15a

The MTX-5-FU-Gem-miR-15a construct was rationally engineered to integrate miRNA-mediated gene regulation with chemotherapeutic functionality and tumor-targeting capability. The guide (antisense) strand of hsa-miR-15a was chemically modified by substituting uridine and cytidine residues with 5-fluorouracil (5-FU) and gemcitabine (Gem), respectively. The passenger (sense) strand was functionalized with a 5′ azide linker and subsequently conjugated with methotrexate (MTX) using dibenzocyclooctyne (DBCO)-mediated copper-free click chemistry. The modified oligonucleotides were synthesized and HPLC-purified (Horizon Discovery). MTX-DBCO was synthesized at Stony Brook University and used for conjugation. Following conjugation, the MTX-linked passenger strand was purified using the Oligo Clean & Concentrator Kit (Zymo Research) according to the manufacturer’s protocol. The purified passenger strand was then annealed with the modified guide strand at 60°C to generate the final duplex construct prior to use in downstream experiments.

### miRNA Transfection

For vehicle-mediated transfection, cells were seeded at a density of 1 × 10⁵ cells per well in 6-well plates and allowed to adhere for 24 hours. Cells were then transfected with 50 nM of the respective miRNA constructs, including MTX-5-FU-Gem-miR-15a, unmodified miR-15a, or scramble control, using Oligofectamine (Thermo Fisher Scientific) according to the manufacturer’s instructions. Five hours following transfection, the media were replaced with fresh media supplemented with 10% dialyzed FBS. For vehicle-free delivery experiments, cells were seeded at 1,500 cells per well in 96-well plates and treated directly with the respective oligonucleotides or drug dilutions in culture media. After 24 hours, the media were replaced with fresh media containing 10% dialyzed FBS to ensure consistent culture conditions.

### 3D Spheroid Culture

Three-dimensional spheroid cultures were established by seeding ovarian cancer cells in ultra-low attachment plates and maintaining them in KnockOut DMEM/F12 medium supplemented with 20% KnockOut Serum Replacement, 1% L-glutamine, 10 ng/mL recombinant human basic fibroblast growth factor (bFGF), and 20 ng/mL recombinant human epidermal growth factor (EGF). Spheroids were allowed to form over a period of 5-7 days under standard culture conditions. For downstream assays, spheroids were dissociated into single-cell suspensions using TrypLE Express at 37°C for 10-15 minutes, followed by gentle mechanical disruption and filtration through a 40 µm cell strainer. Cells were then reseeded in Matrigel-containing medium to allow reformation of spheroids prior to treatment.

### Cytotoxicity Assays

Cytotoxicity of all the miRNAs was assessed without using any delivery vehicle. For cytotoxicity assessment, cells were seeded at a density of 1,500 cells per well in 96-well plates and treated with increasing concentrations of miRNA constructs or drug controls. After six days of treatment, cell viability was measured using the WST-1 Cell Proliferation reagent (MilliporeSigma) according to manufacturer’s protocol. Briefly, 10 µL of WST-1 reagent was added per 100 µL of culture medium and incubated at 37°C for 1 hour. Absorbance was measured at 450 nm with a reference wavelength of 630 nm using a SpectraMax i3 plate reader (Molecular Devices). Relative cell viability was calculated by normalizing absorbance values to those of control-treated cells.

For the Spheroid cells (OVCAR-3/Sph and Sk-OV-3/Sph), the spheroids were dissociated into a single cell suspension and plated at 500 cells/ well in 50μL 10% Matrigel/hCGM in a 96-well plate, black, clear bottom (Falcon, Corning). 72-hours post-plating and visual confirmation of spheroid formation, they were treated with the desired concentrations of either MTX-5-FU-Gem-miR-15a and Olaparib. After 6 days of treatment, cell viability was examined using CellTitre-Glo, 3D (Promega) as per manufacturer’s protocol on plate reader. 100μL of CellTitre-Glo (equivalent to the volume of media in the well) was added to wells, followed by mixing the contents for 2 minutes in an orbital shaker to induce cell lysis. The plate was then incubated at room temperature for 30 minutes, and then luminescence was recorded on the plate reader. Luminescence readings were normalized to the negative control to determine relative cell viability.

Absolute IC50 values were calculated using GraphPad Prism 9 (GraphPad Software, San Diego, CA, USA) using the following equation, where “Y” = 50.

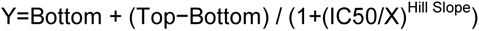

### Cell Cycle Analysis

Cells were seeded in 6-well plates (1 × 10⁵ cells/well) and treated with miRNA constructs or equivalent concentrations of free drugs. After 72 hours, cells were harvested, washed, and resuspended in Krishan buffer containing RNaseH (Thermo Fisher Scientific) and propidium iodide (MilliporeSigma). DNA content was analyzed using a CytoFLEX flow cytometer (Beckman Coulter), and cell cycle distribution was quantified using ModFit LT software (BD Biosciences) to determine the percentage of cells in G₁, S, and G₂ phases.

### Apoptosis Assay

The OC cells were plated and transfected with their respective treatment conditions via vehicle-mediated transfection as described above. 72 h post-transfection, the cells were collected, washed, and stained with Annexin V-FITC (Thermo Fisher Scientific) and propidium iodide (MilliporeSigma) staining. Apoptotic cells were quantified by flow cytometric analysis on a CytoFLEX Flow Cytometer with Annexin V-positive cells categorized as “apoptotic”. Fold change in apoptotic cells was calculated in relation to the number of apoptotic cells observed in the negative control.

### Western Blot Analysis

72 h post-transfection, cells were lysed in RIPA buffer (MilliporeSigma) supplemented with protease inhibitor cocktail (MilliporeSigma) and protein concentrations were determined using a Bradford assay. Equal amounts of protein were separated by SDS-PAGE and transferred to PVDF membranes. Membranes were blocked and incubated with primary antibodies against key targets, including CHK1, WEE1, YAP1, BMI1, BCL-2, BAX, BAK, CCND1, γH2AX, RAD51, p21, GADD45A, cleaved PARP, and cleaved caspase-7. Primary antibodies were diluted in 5% milk (Bio-Rad) in TBST. After staining with primary antibodies, proteins were probed with either secondary antibodies, goat anti-mouse-HRP (Bio-Rad, 1:5000) or goat anti-rabbit-HRP (Bio-Rad, 1:5000), depending on the used primary antibody. Protein bands were visualized using an LI-COR Biosciences Odyssey FC imaging system after the addition of SuperSignal West Pico PLUS Chemiluminescent Substrate (Thermo Fisher Scientific). Protein expression was quantified by densitometric analysis of band intensities using Image Studio software (version 5.2.4; LI-COR Biosciences). For each target protein, expression levels were normalized to β-actin and reported relative to the negative control sample, which was set to 1.

### RNA Sequencing and Bioinformatic Analysis

OVCAR-3 parental (PAR) and olaparib-resistant (OR) cells were transfected as described above, and total RNA was isolated 72 hours post-transfection using TRIzol reagent (Thermo Fisher Scientific) according to the manufacturer’s instructions. RNA samples were submitted to Novogene for poly(A)-enriched mRNA sequencing. Raw sequencing data were processed by Novogene, and normalized gene expression and differential expression outputs were provided for downstream analysis.

Subsequent analyses were performed using R (v4.2) and Python (v3.9). In R, gene set enrichment analysis (GSEA) was conducted using the clusterProfiler package, with genes ranked based on log2 fold-change values for each comparison (PAR and OR conditions). GSEA was performed using KEGG pathway gene sets with the gseKEGG function, based on the original GSEA framework and implemented using a fast algorithm ^73,74^. Functional enrichment analyses were conducted using KEGG pathways and Gene Ontology (GO) annotations ^75,76^.

Python was used for additional data processing, integration, and visualization of transcriptomic datasets. For integrated analyses, a merged expression matrix comprising PAR NC, PAR M15a, OR NC, and OR M15a conditions was log2-transformed and used for principal component analysis (PCA) based on the top variable genes. Heatmaps were generated using row-wise Z-score normalization of selected gene sets.

A pathway “reversion” score was defined as:

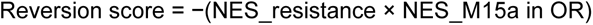

where NES_resistance represents the normalized enrichment score obtained from the comparison of untreated olaparib-resistant cells versus untreated parental cells (OR NC vs PAR NC), reflecting resistance-associated pathway enrichment. NES_M15a in OR represents the normalized enrichment score from the comparison of treated versus untreated olaparib-resistant cells (OR M15a vs OR NC), capturing treatment-induced pathway modulation in the resistant state. Pathways related to DNA damage response, replication stress, cell cycle regulation, apoptotic signaling, and oncogenic survival pathways were prioritized for analysis and visualization.

All analyses and visualizations were performed using R and Python, and scripts are available upon request.

### Combination and Synergy Analysis

The combinatorial effects of MTX-5-FU-Gem-miR-15a and olaparib were evaluated using a matrix-based dose-response approach. Ovarian cancer cells were seeded in 96-well plates at a density of 1,500 cells per well and allowed to adhere for 24 hours under standard culture conditions. Following attachment, cells were treated with combinations of MTX-5-FU-Gem-miR-15a and olaparib arranged in a 8 × 6 dose matrix, in which increasing concentrations of MTX-5-FU-Gem-miR-15a were combined with increasing concentrations of olaparib across columns and rows, respectively. For MTX-5-FU-Gem-miR-15a, concentrations ranged from 0 nM to 5 nM, while olaparib concentrations ranged from 0 µM to 5-10 µM, selected based on prior single-agent IC₅₀ values to capture sub-effective, intermediate, and effective dose ranges for both agents.

Cells were incubated with the combination treatments for 6 days, consistent with single-agent cytotoxicity assays. At the endpoint, cell viability was assessed using the WST-1 Cell Proliferation Assay, and absorbance was measured at 450 nm with a reference wavelength of 630 nm. Raw absorbance values were normalized to untreated controls to determine relative cell viability across all treatment conditions.

Drug interaction effects were quantified using the Bliss independence model implemented in the SynergyFinder platform. Bliss synergy scores were calculated by comparing the observed combined effect to the expected additive effect of each agent based on single-agent responses. Synergy scores were interpreted using established thresholds, where a Bliss score greater than 10 was considered synergistic, values between 0 and 10 were considered additive, and values below 0 were considered antagonistic. Global synergy scores were calculated across the entire dose-response matrix, and synergy heatmaps were generated to visualize interaction patterns between MTX-5-FU-Gem-miR-15a and olaparib across all tested concentration combinations.

### Metastatic Ovarian Cancer Mouse Model

All *in vivo* experiments were conducted in compliance with the Stony Brook University Institutional Animal Care and Use Committee (IACUC) guidelines. Non-obese diabetic (NOD)/severe combined immunodeficiency (SCID) mice (JAX: 001303) were purchased from The Jackson Laboratory (The Jackson Laboratory). To establish a metastatic model, 8-week-old female NOD/SCID mice (n = 7-8 per group) were inoculated with 1.5 x 10^6^ SK-OV-3 (+Luc) cells suspended in 0.2 mL of PBS via intravenous (IV) tail vein injection to establish a metastatic model. One-week post-inoculation, metastatic tumor formation was confirmed by bioluminescent IVIS imaging. Animals were randomized into two groups and treated with either 75 μg of PEI-MAX (Polysciences) in 5% D-glucose (vehicle control group) (Thermo Fisher Scientific) or 75 μg (3.75 mg/kg) of MTX-5-FU-Gem-miR-15a miRNA with 75 μg of PEI-MAX in 5% D-glucose (treated group). All the treatments were administered via intravenous tail vein injection on alternating days for 2 weeks (a total of 8 doses).

The same experimental design was repeated using luciferase-expressing SK-OV-3/OlaR cells, with animals assigned to three groups: vehicle control, IV-treated, and intraperitoneally (IP)-treated. Treatment doses were identical to those used in the parental SK-OV-3 model.

Luciferase expression was used to measure tumor growth using the IVIS Spectrum *in vivo* Imaging System (IVIS) (PerkinElmer) as previously described. In brief, mice were injected with 100 ul of IVISBrite D-Luciferin, RediJect (PerkinElmer). 10 min post-injection, *in vivo* luciferase expression was measured via IVIS. Tumor growth was calculated as a function of (total flux at time of measurement [p/s])/(total flux at initial measurement [p/s]).

Luciferase-expressing ovarian cancer cells were implanted into immunocompromised mice, and tumor growth was monitored longitudinally using IVIS imaging. Animals were treated with MTX-5-FU-Gem-miR-15a, and tumor burden was quantified over time based on bioluminescence signal intensity.

## QUANTIFICATION AND STATISTICAL ANALYSIS

All experiments were performed in biological triplicates unless otherwise specified and data were analyzed using GraphPad Prism 9 software (GraphPad Software, San Diego, CA, USA). Statistical comparisons between two groups were conducted using an unpaired Student’s T-test. For *in vivo* studies, tumor growth kinetics and treatment effects over time were analyzed using two-way ANOVA with mixed-effects modeling to account for repeated measurements across time points. Data are presented as mean ± standard deviation (SD). Statistical significance was defined as a p-value < 0.05. Levels of significance are indicated as follows: p < 0.05 (*), p < 0.01 (**), p < 0.001 (***), and p < 0.0001 (****).

