## Supplemental figures for "Developing a Multimodal miR-15a Mimic to Overcome PARP Inhibitor Resistance in Epithelial Ovarian Cancer"

### Figure S1. Related to Figure 1

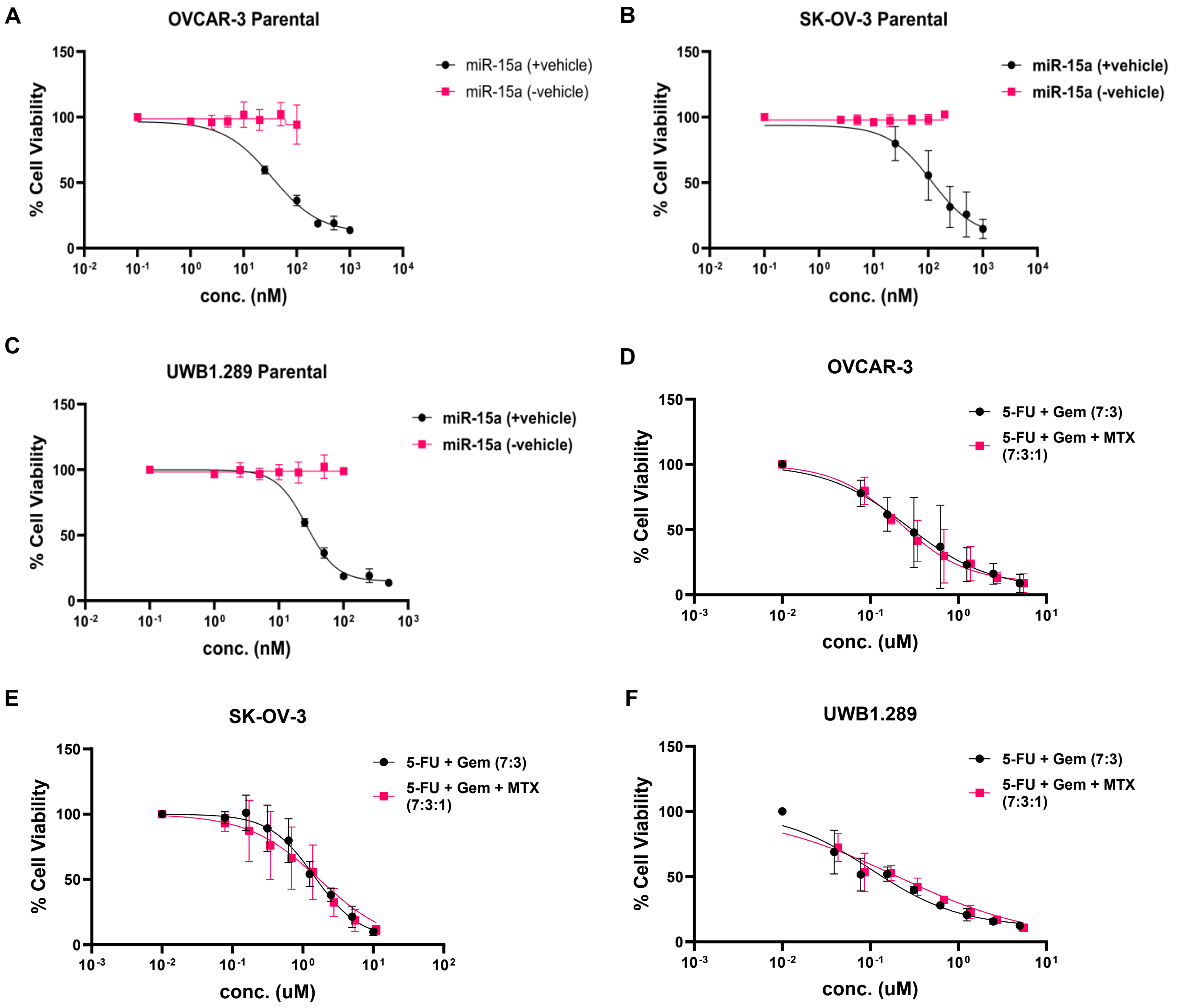


**Figure S1. Vehicle dependency of unmodified miR-15a and limited cytotoxicity of corresponding drug combinations**

(A-C) Dose–response curves of unmodified miR-15a in the presence (+vehicle) or absence (-vehicle) of transfection reagent in OVCAR-3 (A), SK-OV-3 (B), and UWB1.289 (C) cells. Cells were treated with increasing concentrations of miR-15a in presence or absence of Oligofectamine, and cell viability was measured by WST-1 assay after 6 days.

(D-F) Dose–response curves of chemotherapeutic combinations corresponding to the modified construct in OVCAR-3 (D), SK-OV-3 (E), and UWB1.289 (F) cells. Cells were treated with 5-FU + gemcitabine (7:3) or 5-FU + gemcitabine + methotrexate (7:3:1) for 6 days, followed by WST-1 assay.

Data are presented as mean ± SD.

### Figure S2. Related to Figure 4

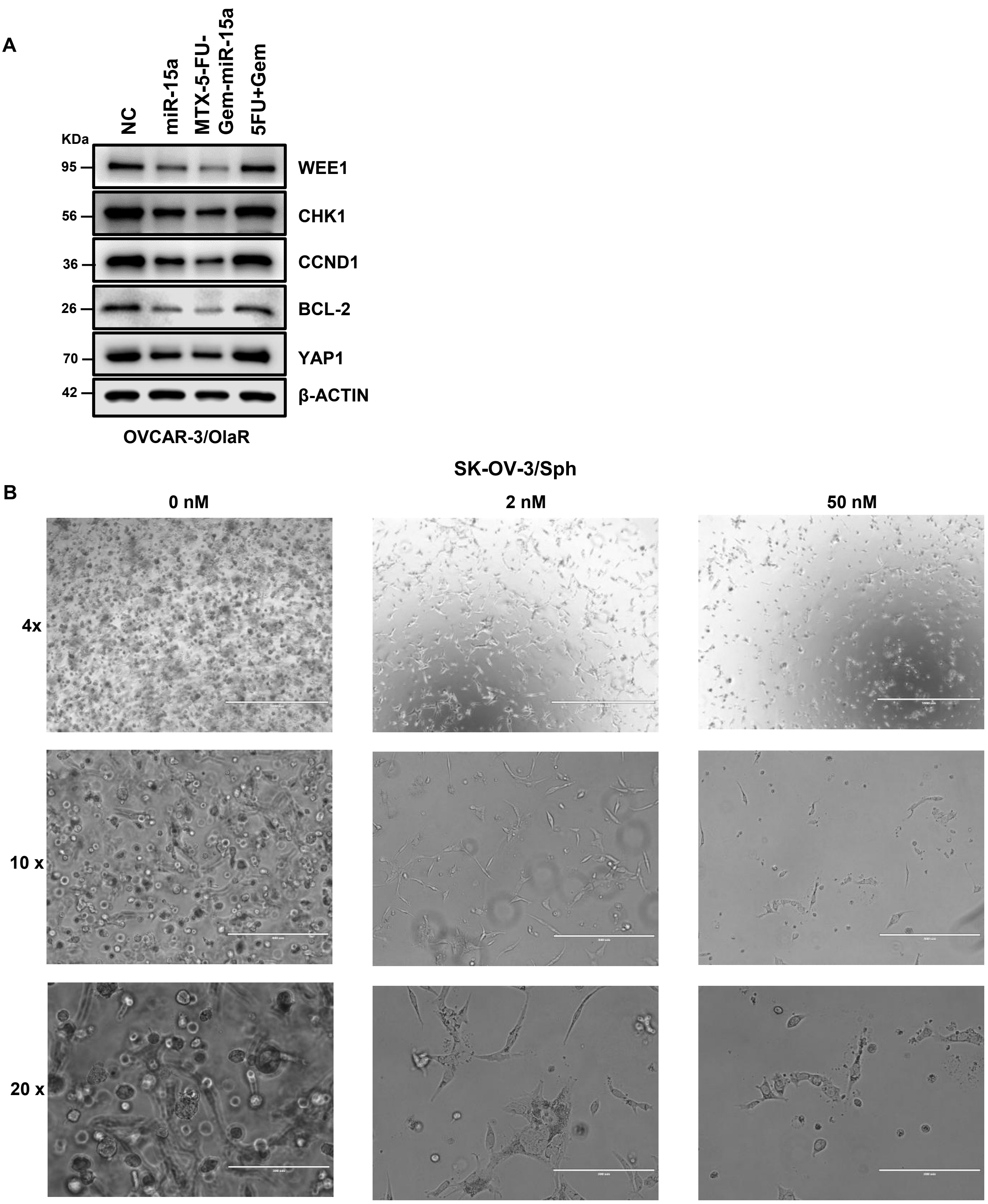


**Figure S2. MTX-5-FU-Gem-miR-15a retains target specificity and induces dose-dependent cytotoxicity in olaparib-resistant and 3D spheroid models.**

(A) Western blot analysis of miR-15a target proteins in olaparib-resistant ovarian cancer cells following treatment with MTX-5-FU-Gem-miR-15a, demonstrating retention of target specificity in olaparib resistant cells.

(B) Representative brightfield images of SK-OV-3 – derived spheroids treated with increasing concentrations of MTX-5-FU-Gem-miR-15a, showing dose-dependent disruption of spheroid integrity. Higher-magnification images of spheroids under corresponding treatment conditions, highlighting cellular disaggregation and morphological changes associated with cytotoxic response. Scale bars are indicated.

**Figure S3. Related to Figure 5**

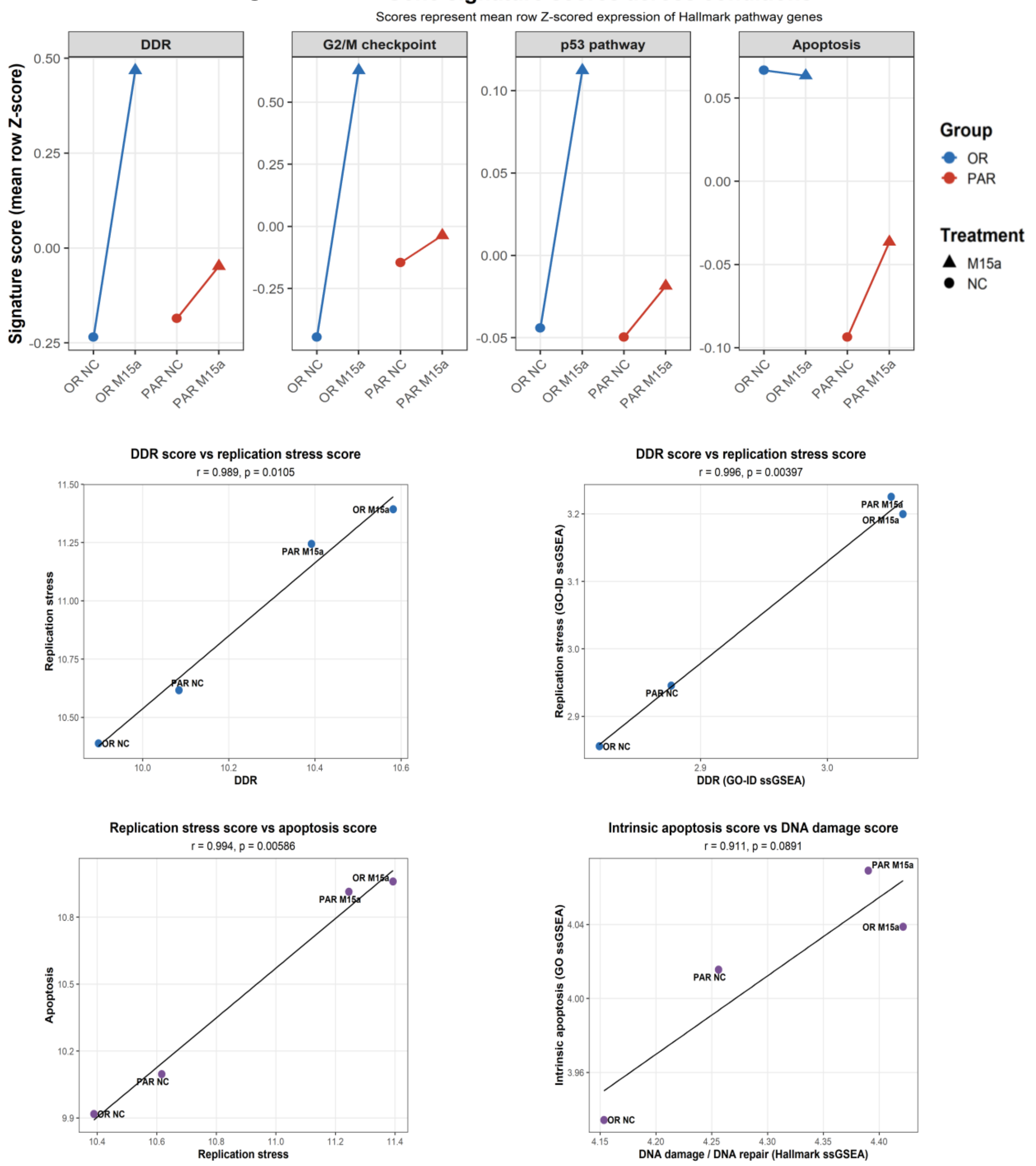


**Figure S3. MTX-5-FU-Gem-miR-15a induces coordinated activation of DNA damage, replication stress, and apoptotic signaling pathways**

(A) Gene signature scores for selected Hallmark pathways, including DNA damage response (DDR), G2/M checkpoint, p53 signaling, and apoptosis, in parental (PAR) and olaparib-resistant (OR) OVCAR-3 cells treated with MTX-5-FU-Gem-miR-15a (M15a) or negative control (NC). Scores represent mean row Z-scored expression values of genes within each pathway.

(B-C) Correlation between DDR and replication stress scores derived from Hallmark gene sets (B) and Gene Ontology (GO) gene sets (C).

(D) Correlation between replication stress and apoptosis scores derived from Hallmark gene sets.

(E) Correlation between intrinsic apoptotic signaling and DNA damage/repair scores derived from Gene Ontology (GO) gene sets.

Pearson correlation coefficients (r) and corresponding p values are indicated in each plot.

### Figure S4. Related to Figure 7

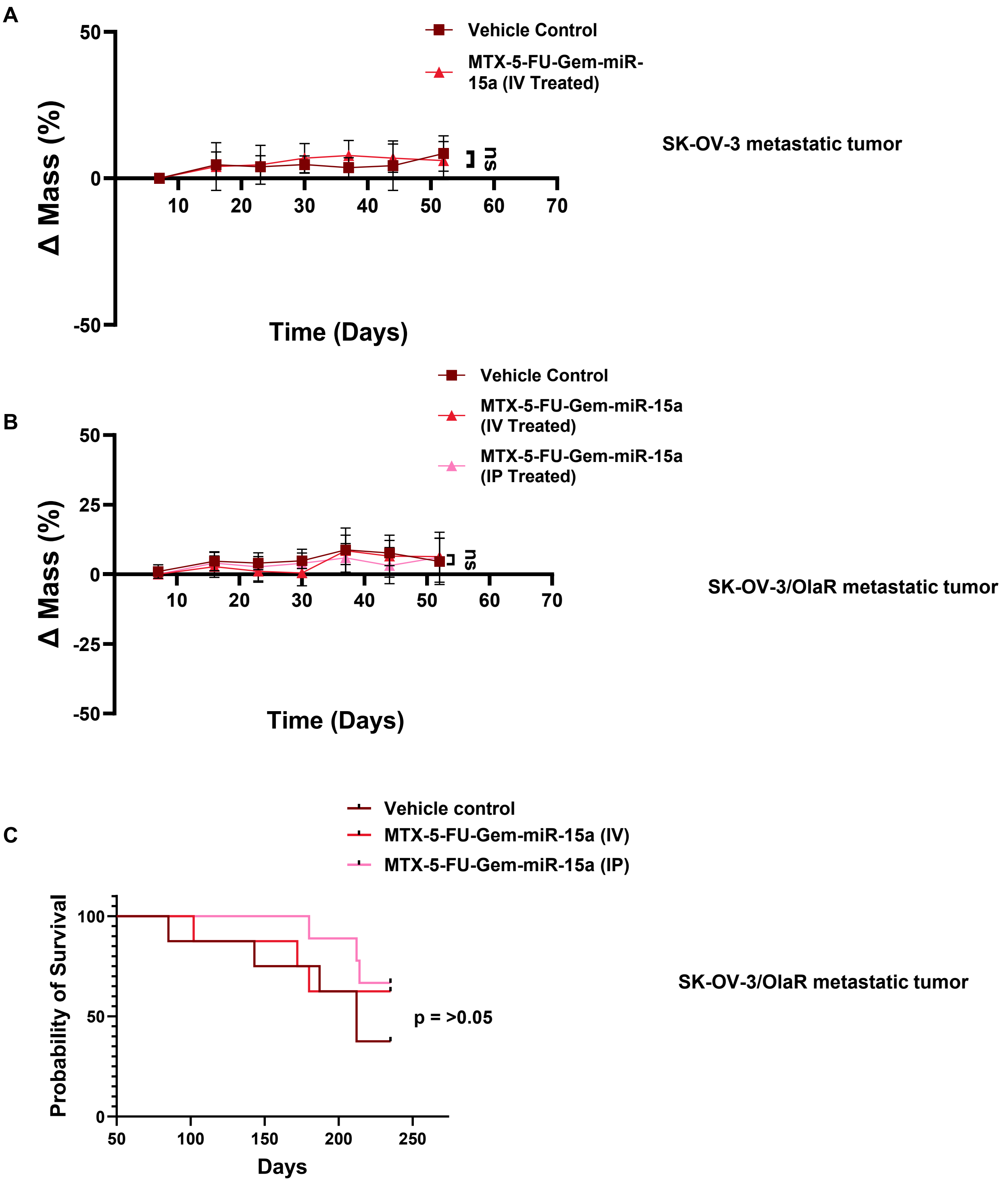


**Figure S4. MTX-5-FU-Gem-miR-15a is well tolerated and improves survival in parental and resistant SK-OV-3 models.**

(A) Body weight changes over time in the SK-OV-3 metastatic tumor model under the indicated treatment conditions. Body weight was monitored at the indicated time points throughout the treatment period.

(B) Body weight changes over time in the SK-OV-3/OlaR tumor model under the indicated treatment conditions. Measurements were recorded at each time point during the study.

(C) Kaplan-Meier survival curves of mice bearing SK-OV-3/OlaR tumors under the indicated treatment conditions. Survival was monitored over the course of the study and analyzed using the Kaplan-Meier method.

Data in (A) and (B) are presented as mean ± SD.

| Cell line | Hsa-miR-15a (-Vehicle) | Hsa-miR-15a (+Vehicle) | 5-FU + Gem (7:3) | 5-FU + Gem + MTX (7:3:1) |
|---|---|---|---|---|
| **OVCAR-3** | na | 40 nM | 1454 nM | 1353 nM |
| **SK-OV-3** | na | 115 nM | 266 nM | 220 nM |
| **UWB1.289** | na | 26 nM | 133 nM | 161 nM |

**Table S1. IC50 values of miR-15a and drug combinations across ovarian cancer cell lines. Related to Figure S1.**
IC50 values were determined from dose-response curves generated using WST-1 assays after 6 days of treatment. Values are reported in nM. “na” indicates not applicable.
